# Identity-by-descent captures shared environmental factors at biobank scale

**DOI:** 10.1101/2025.05.03.652048

**Authors:** Franco Marsico, Silvia Buonaiuto, Ernestine K Amos-Abanyie, Lokesh K Chinthala, Akram Mohammed, Rima Zahr, Regeneron Genetics Center, Robert J Rooney, Robert W Williams, Robert L Davis, Terri H Finkel, Chester W Brown, Marcella Vacca, Pjotr Prins, Vincenza Colonna

## Abstract

Genealogical ties leave traces beyond shared genetic material: relatives tend to share environmental exposures and cultural practices that persist across generations. Using genomic and electronic health record data from 13,143 individuals in the Biorepository for Integrative Genomics, we applied a hierarchical community detection algorithm to group individuals into four major communities and 17 subcommunities based on the proportion of their genome shared identical by descent (IBD). This approach captured a fine-scale demographic structure beyond traditional ancestry classifications. By integrating neighborhood-level geographic data with census-derived environmental metrics, we revealed unequal exposure to environmental stressors between subcommunities, which correlated with differential rates of health conditions, including elevated respiratory condition prevalence in subcommunities under greater pollution burden. Through an interventional mediation analysis we found that excess risk at the subcommunity-level substantially exceeded what measured environmental factors alone could explain. thus showing the contribution of unmeasured shared exposures that IBD-based grouping captures. Subcommunity-level disparities persisted in sensitivity analyses adjusting for self-reported race, indicating that IBD-based clustering captures health-relevant information beyond demographic categories commonly used as proxies for social determinants of health. We implemented an open-source dashboard for interactive exploration of IBD-defined communities, disease prevalence, and environmental exposures. Together, we demonstrate that IBD-based clustering jointly captures genetic and environmental determinants of health, offering a scalable framework for precision health and population genetics that translates biobank data into actionable insights.

## 1 Introduction

Genomic studies increasingly rely on large biobanks that link genetic, clinical, and environmental data (Buonaiuto et al., 2025; Belbin et al., 2021; Nait Saada et al., 2020; Caggiano et al., 2023). How participants in these resources are grouped shapes which patterns of variation become detectable. Identity-by-descent (IBD) supports a grouping based on the genealogical relationships among individuals, inferred directly from the genome.

IBD refers to DNA segments shared by two individuals due to inheritance from a common ancestor (Palamara et al., 2012). Recombination progressively fragments these segments across generations, so the total amount of IBD shared decreases with genealogical distance. Close relatives, first and second degree, usually share more than just genetic material: their relationships are the foundation of many family systems (Itao and Kaneko, 2021) and typically reflect shared culture, social norms, and environments. These ties extend beyond close kin: even distant relatives can share cultural traits passed through generations of slowly evolving social norms, as observed by Cavalli-Sforza and Feldman (1981; 1986).

To study population structure, the common practice of categorizing individuals into discrete genetic groups faces challenges that go beyond technical limitations. Such categorizations impose artificial boundaries on what is fundamentally a continuous spectrum of human genetic variation (Barbujani and Colonna, 2010; Dobzhansky, 1962), and depend on predefined reference populations whose composition is itself the product of sampling decisions (Rosenberg et al., 2002; Barbujani et al., 1997; Maples et al., 2013). IBD-based grouping addresses these limitations by recovering population structure directly from genealogical relationships among participants, without recourse to external reference panels (Browning et al., 2018; Browning and Browning, 2011; Ralph and Coop, 2013; Caggiano et al., 2023; Gilbert et al., 2022; Naseri et al., 2021; Dai et al., 2020; Nait Saada et al., 2020; Belbin et al., 2021; Shemirani et al., 2026). Recent consensus statements from the National Academies of Sciences and editors of biomedical journals converge in this direction, calling for the differentiation of genetic ancestry from race and ethnicity and supporting reference-free frameworks where possible (National Academies of Sciences, Engineering, and Medicine, 2023; Lewis et al., 2024). Self-reported race and ethnicity categories retain practical value in U.S. clinical settings, where researchers often argue they function as proxies for social and environmental factors such as residential context and access to care (Manski et al., 2023).

In this study, we used data from the Biorepository for Integrative Genomics (BIG) cohort (Buonaiuto et al., 2025) to build IBD-based communities that integrate genomic, clinical, and environmental information, providing a scalable framework to study health disparities and the co-segregation of genealogy and environment at biobank scale. The BIG resource provides comprehensive electronic health records (EHR), neighborhood-level environmental data linked through ZIP codes, and genomic information from 13,143 participants, with particular emphasis on pediatric populations. A key feature of BIG is its representation of high genetic diversity concentrated within a relatively confined geographic area, allowing the investigation of the interplay between genetics, the environment and health using standardized data.

Here, we demonstrate that IBD communities are consistent with inferred and self-reported ancestry while capturing a fine-scale genetic substructure with distinct environmental exposures and health outcomes, reflecting geographic stratification at the neighborhood level. We show that community-specific health patterns can be traced to differential environmental exposures, illustrating how genealogical relatedness and shared environment co-segregate to shape disease risk at the population level. Mediation analyses indicate that measured environmental factors explain only part of the observed health disparities, with a residual covariation with IBD-defined community membership that likely reflects unmeasured shared exposures. Our findings establish IBD communities as a valuable tool for precision public health, enabling targeted interventions based on community-specific genetic and environmental profiles. Together, these findings demonstrate that the genealogical structure inferred from genomic data captures health-relevant variation beyond conventional demographic categories, providing a scalable framework to study population health and the co-segregation of genealogy and environment in biobank-scale datasets.

## 2 Results

### 2.1 Capturing population structure using information from genomic segments shared identical by descent

The BIG initiative has enrolled over 42,000 participants with linked EHRs and collected more than 15,000 biosamples (**Supplementary Figure S1**). The cohort is predominantly pediatric (87% under 18 years), spanning ages from infancy to 90 years (mean = 8.4, median = 6.2) (Buonaiuto et al., 2025). Unlike typical pediatric cohorts of healthy mother–child pairs, BIG focuses on children with diverse diseases and ancestry backgrounds.

To assess population structure, we performed identity-by-descent (IBD) analysis, which infers genetic relationships directly from shared genomic segments (Zhou et al., 2020). Total pairwise IBD was calculated as the sum of the length (cM) of all shared segments between individuals. The distribution of total IBD values followed an exponential decay (**Figure 1A**), indicating that distant relatives pairs far outnumber close kinships, consistent with previous studies (Caggiano et al., 2023; Nait Saada et al., 2020; Shemirani et al., 2021).

**Figure 1:**
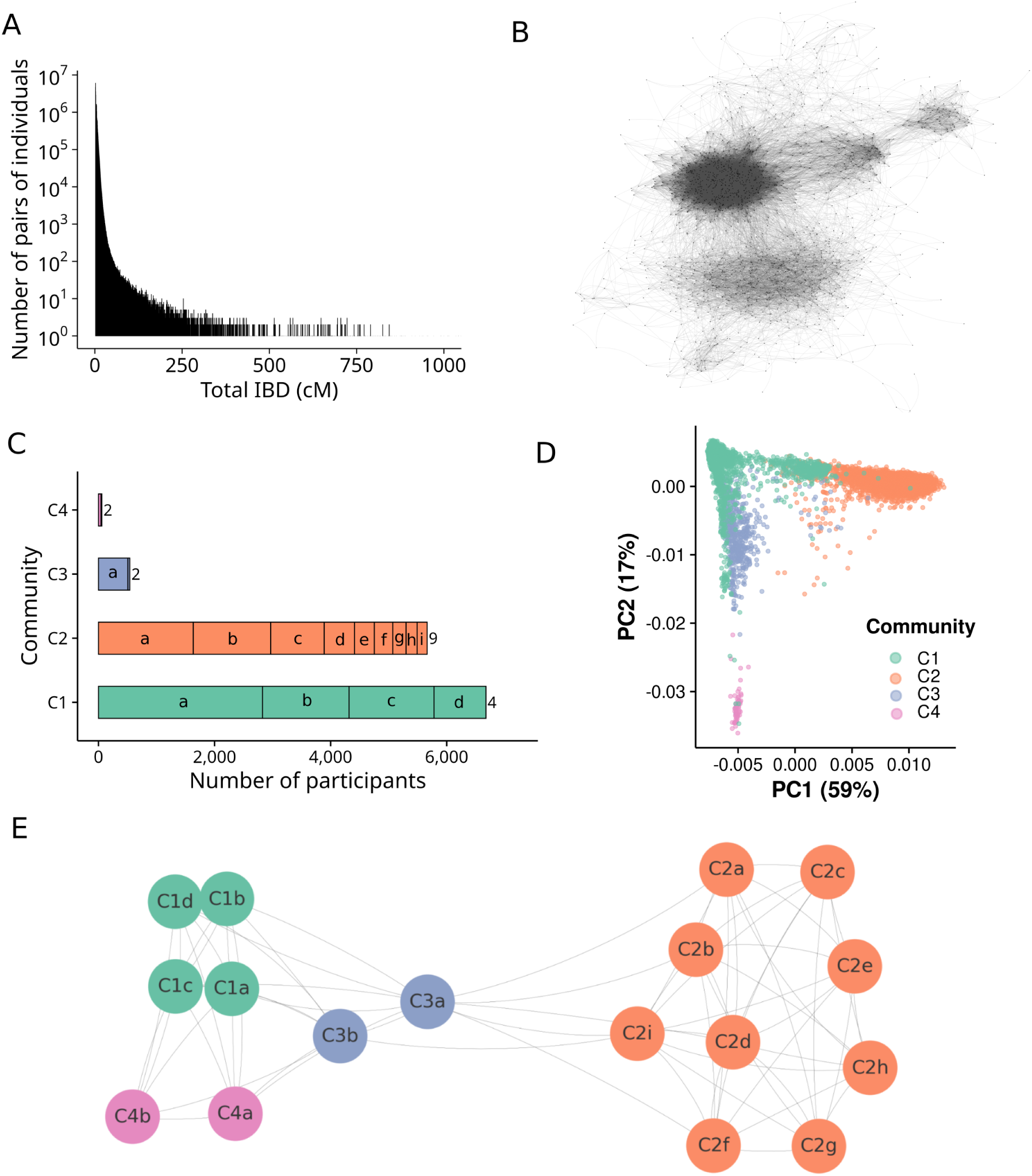
Identity-by-descent (IBD) community detection on the BIG cohort. **(A)** Exponential decay of total IBD (cM). **(B)** IBD-based network with 13,143 nodes, revealing four denser areas surrounded by sparse zones, suggesting a structured network. Fruchterman-Reingold layout was used to do the plot (Gajdoš et al., 2016). **(C)** Community sizes after the first Leiden iteration (C1 to C4), and size of the subcommunities (small case letters on bars) derived from the second Leiden iteration. **(D)** PCA plot showing samples distribution across the two first PCs. Colors represent the communities. The first two principal components explain 76% of the variance captured by the first 20 PCs. **(E)** Network of subcommunities highlighting diversity and inter-connectivity (Fruchterman-Reingold layout). Connections between subcommunities are based on total IBD.

We constructed an IBD network, where nodes represent individuals (n = 13,143) and edge lengths are inversely proportional to shared IBD. The network’s global density (0.018) and average degree (241.15) indicate a well-connected structure. Four main dense clusters are visible, surrounded by smaller peripheral groups (**Figure 1B**). Using the Leiden algorithm (Anuar et al., 2021), we applied two rounds of hierarchical community detection. The first iteration identified four major communities (C1–C4), two of which contained more than 5,000 individuals. Subsequent iterations within each community revealed 17 subcommunities (**Figure 1C**).

The community classification corresponds well with the population structure revealed by PCA of autosomal genetic markers, with clear separation between communities (**Figure 1D**). This concordance between discrete community assignments and continuous genetic variation, both inferred from the same genomic data, is consistent with communities reflecting non-arbitrary genealogical structure rather than artificial partitions. Despite this overall trend, we also notice individuals positioned between the main groups in the PCA space, as well as some individuals from one community who fall within the region predominantly occupied by a different community. As an example, C1 individuals in C2 space would be assigned to C2 based on their position in the PCA-based classification; however, the IBD-based approach places them in C1 due to detected signals of relatedness (up to third degree), highlighting the additional power of relationship-based community detection over purely dimensional-reduction-based methods.

At a higher level, a network of subcommunities (**Figure 1E**) shows that the number of subgroups does not correlate with parent community size, suggesting that internal genetic diversity influences subdivision. Community C2 exhibits the highest genetic diversity and the greatest number of subcommunities, supported by pairwise Fst values (**Supplementary Figure S2**). In contrast, C1 subcommunities are more closely connected, indicating closer relationships among members, while C3 connects to both C1 and C2, probably reflecting shared ancestry patterns.

### 2.2 IBD communities reflect geographical ancestry with fine-scale substructure

Having determined IBD-based communities, we next asked whether they corresponded to global ancestry previously inferred from reference populations (Buonaiuto et al., 2025). We found enrichment of EUR-inferred ancestry (91 %) in C1, AFR-inferred (49 %) with EUR–AFR admixed components (39 %) in C2, EUR–AMR admixed (51 %) with multi-component admixture (43 %) in C3, and EAS-inferred (81%) in C4 (**Figure 2A**). Community and ancestry, both inferred from the same genomic data, showed strong concordance (Cramér’s V = 0.81).

**Figure 2:**
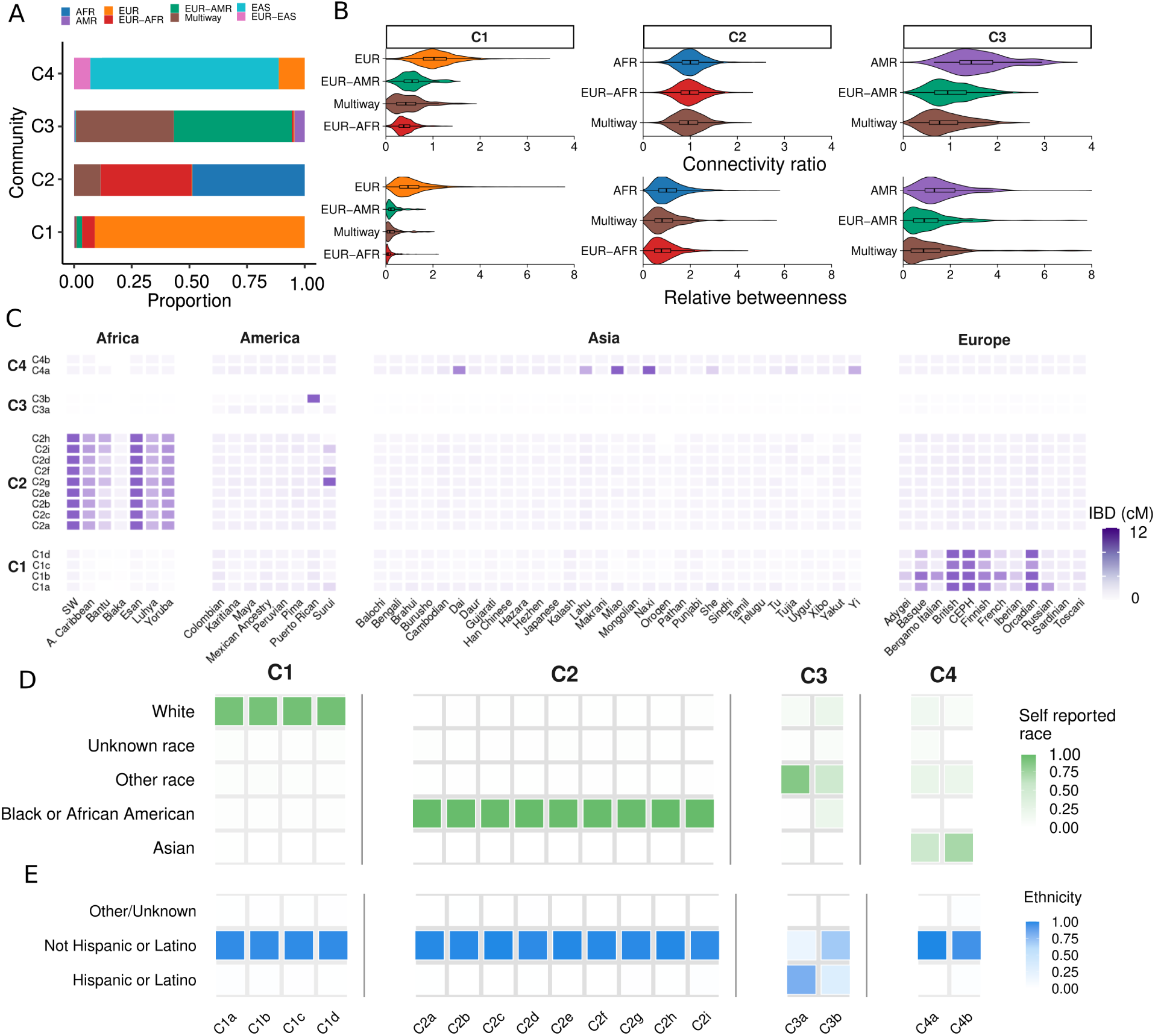
Genetic and demographic patterns across sub-communities. **(A)** Inferred continental and admixed ancestries per community from previous work. **(B)** Ancestry’s role in the IBD-network. Connectivity ratio for each node was defined as its degree divided by the mean degree of its IBD community. Values greater than 1 indicate higher connectivity compared to the average. Betweenness measures the centrality of individuals in communities; individuals with betweenness above 1 are more central, while values below 1 indicate more peripheral individuals. **(C)** Heatmap showing mean IBD values between subcommunities and reference populations, highlighting distinct genetic relationships across C1 to C4. **(D)** Distribution of self-reported race (HL7 v2.5 categories), and **(E)** self-reported Hispanic/Latino ethnicity proportions within subcommunities.

We characterized the relationship between ancestry and network topology by computing each BIG individual’s connectivity (degree link strength) and betweenness (path-based centrality). Stratification by ancestry revealed that individuals with predominantly EUR-inferred ancestry in C1, predominantly AFR-inferred in C2, and predominantly AMR-inferred in C3 tend to act as hub–bridge nodes, mediating many connections and shortest paths (**Figure 2B**). In contrast, individuals with multi-component ancestry inference exhibit lower connectivity and betweenness, occupying more peripheral network positions. Those with multi-way (≥3 components) ancestry profiles consistently fall into low-degree, low-betweenness niches, while certain moderately connected nodes display elevated betweenness, acting as critical inter-community bridges. C4 communities were not analyzed due to their smaller size.

To further characterize communities and subcommunities, we measured IBD sharing between BIG subcommunities and the HGDP-1kGP reference panels. The IBD sharing patterns align with the ancestry correlations we previously inferred (**Figure 2C**). However, the fine-scale subcommunity structure suggests subtle demographic events that shaped genetic diversity within broader continental groups. For example, subcommunity C3b showed a stronger relationship with the Puerto Rican population, characterized by its three-way ancestry pattern (Browning et al., 2018; Mas-Sandoval et al., 2023), while C3a was more closely associated with Peruvian and other AMR populations. This subdivision suggests multiple waves or routes of population movement and mixing within C3. Likewise, the connections within C2 (predominantly enriched in AFR-inferred ancestry) of C2g with Surui (HGDP population from America superpopulation) might indicate shared ancestry components that transcend simple continental divisions, possibly reflecting more recent demographic events.

After characterizing community structure through genetic measures, we next examined how communities aligned with participants’ self-reported race and ethnicity (**Figure 2D-E**). Similar to what was observed for IBD sharing with reference populations, while community-level patterns aligned with self-reported categories as expected for an urban U.S. cohort with documented residential segregation history (C1: 91% white, C2: 97% Black/African American, C4: 62% Asian), subcommunity analysis reveals important nuances. Most notably, C3 subcommunities diverge markedly. In terms of self-reported race, C3a is predominantly Other race (73%), whereas C3b shows a three-way split across Other race (33%), White (21%), and Black/African American (21%). In terms of self-reported ethnicity, individuals in C3a were more likely to self-identify as Hispanic or Latino (72.3%), while those in C3b predominantly self-report as Not Hispanic or Latino (69.6%).

Overall, our findings demonstrate that IBD-based communities capture population stratification consistent with both genetic ancestry and self-reported race/ethnicity. While major communities reflect continental ancestry patterns, subcommunity analysis reveals finer population substructure and demographic complexity, including admixture patterns and the nuanced relationships between genetic ancestry and self-identity. This hierarchical approach provides the resolution to understand both historical demographic processes and contemporary genetic diversity within the BIG cohort.

### 2.3 IBD sharing reflects shared environments in the BIG cohort

To elucidate the spatial distribution of genetic relationships, we examined whether subcommunities exhibit distinct geographical patterns and how these relate to IBD. Specifically, we geolocated BIG participants’ ZIP codes at the neighborhood level using publicly available spatial polygons (Pebesma and Bivand, 2018), and then analyzed whether genetically determined subcommunities cluster geographically. We detected a strong association between genealogical and geographical distances (exponential fit *R*^2^ = 0.969, p-value *<* 0.0001, **Figure 3A**, **Supplementary Figure S3**). Nearly half (49%) of close relatives (first- and second-degree) lived in the same neighborhood, while the remaining pairs typically resided 4-6 km apart. Third-and fourth-degree relatives showed greater separation, though still within 20 km, with 34.3% sharing the same neighborhood. Beyond the seventh or eighth degree, geographic distance increased sharply, only 4.45% of eighth-degree or more distant relatives lived in the same neighborhood.

**Figure 3:**
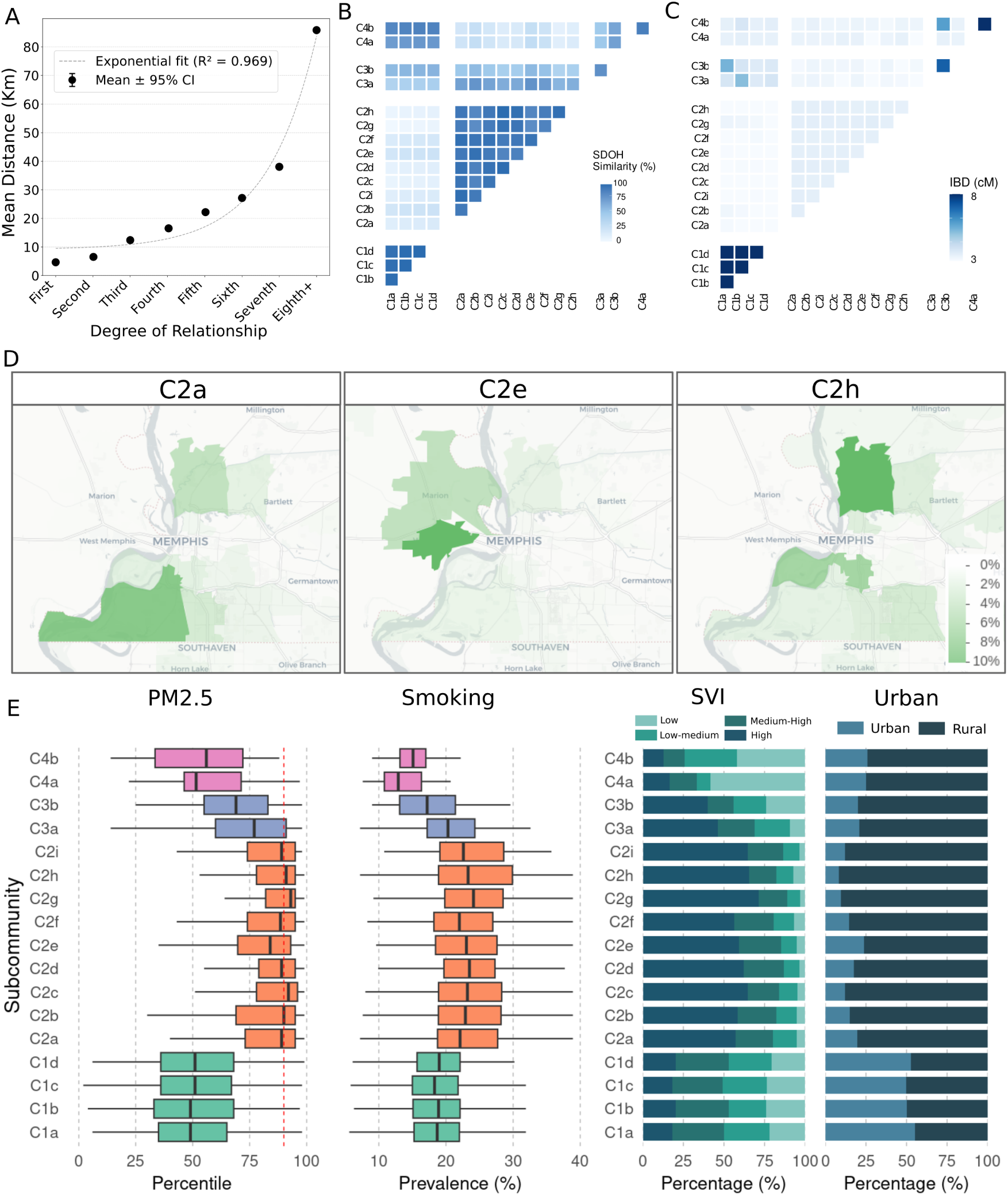
Genealogy, environment and geography of BIG communities. **(A)** Relationship between genealogical (degree of relationship) and geographical (Km) distances. Exponential fit is shown. **(B)** Social determinants of health similarity index at neighborhood level across communities. The diagonal of the matrix is not shown and corresponds to the maximum (100%) identity. **(C)** IBD sharing measured as median pairwise IBD (cM) between pairs of different subcommunities **(D)** Maps centered on the city of Memphis showing the geographical distribution of individuals from three subcommunities (C2a, C2e and C2h) from community C2. **(E)** Social determinants of health by subcommunities. PM_2.5_ EJ index is an environmental justice metric derived from the ambient concentration of fine particulate matter, red line indicates the 90th percentile, above which indicates high risk. Smoking prevalence, social vulnerability index and urbanization are also reported.

We used ZIP codes to link communities to social determinants of health (SDoH), i.e., non-medical factors influencing health outcomes. All SDoH values were obtained from publicly available datasets. Complete results, including interactive visualizations and detailed methodology, are available on our dedicated website (https://github.com/MarsicoFL/BIG.Communities). We quantified SDoH similarity across subcommunities using mean SDoH and found that subcommunities within the same community shared highly similar social environments, while communities differed markedly from each other (**Figure 3B**). This pattern held even for C1, whose subcommunities maintained 96% SDoH similarity despite being the most geographically dispersed (median distance 88.6 km between individuals vs. 18.9-22.1 km for C2 and C3, **Supplementary Figure S4B**). Conversely, C2 exhibited the greatest social heterogeneity among its subcommunities while occupying a smaller geographic area than C1 (**Supplementary Figure S4B**). SDoH similarity is strongly correlated with mean IBD sharing within and between subcommunities. To account for non-independent pairwise entries, we tested the association using a Mantel permutation. We find a monotonic association (Mantel Spearman *ρ* = 0.747, *p* = 2.0 × 10*^−^*^4^), demonstrating that communities with similar SDoH share greater genetic relatedness (**Figure 3C**). In addition, the nuanced variation in SDoH within C2 is reflected in overall lower IBD sharing within C2. The spatial separation of C2 subcommunities likely explains their SDoH variation. For example, subcommunities C2a, C2e, and C2h occupy distinct western, central, and eastern regions of Memphis respectively (**Figure 3D**, Supplementary Figure S4A), demonstrating how IBD-based clustering captures the interplay between genetic relatedness, geographic separation, and differential social environments.

Large differences between communities and more subtle differences between subcommunities also appear when examining in more detail key SDoH factors with direct impacts on clinical outcomes. As shown in **Figure 3E**, subcommunities within group C2 exhibit elevated PM_2.5_ Environmental Justice (EJ) index values, indicative of higher pollution levels, coupled with increased urban residency and smoking prevalence. Moreover, the Social Vulnerability Index reveals that a substantial fraction of C2 resides in high-risk areas, a trend that is also seen, though less pronounced, in C3.

These findings suggest that in the BIG cohort IBD-based community membership strongly reflects social environmental exposure. IBD-based communities likely represent distinct socioeconomic or cultural groups with differential access to resources and environmental conditions, which may have important implications for understanding health disparities across these populations.

### 2.4 Health condition patterns in subcommunities reflect differential environmental exposure

Having assessed the differential environmental exposures across communities and subcommunities, we next examined whether these differences translate into disparities in disease incidence. We mapped the ICD-9/10 codes from the BIG electronic health records to PheCode categories and found overall distinct disease prevalence patterns across communities (**Figure 4B**). Community-level analysis revealed differences in environmentally-influenced conditions (Tran et al., 2023; Eisenberg et al., 2007).

**Figure 4:**
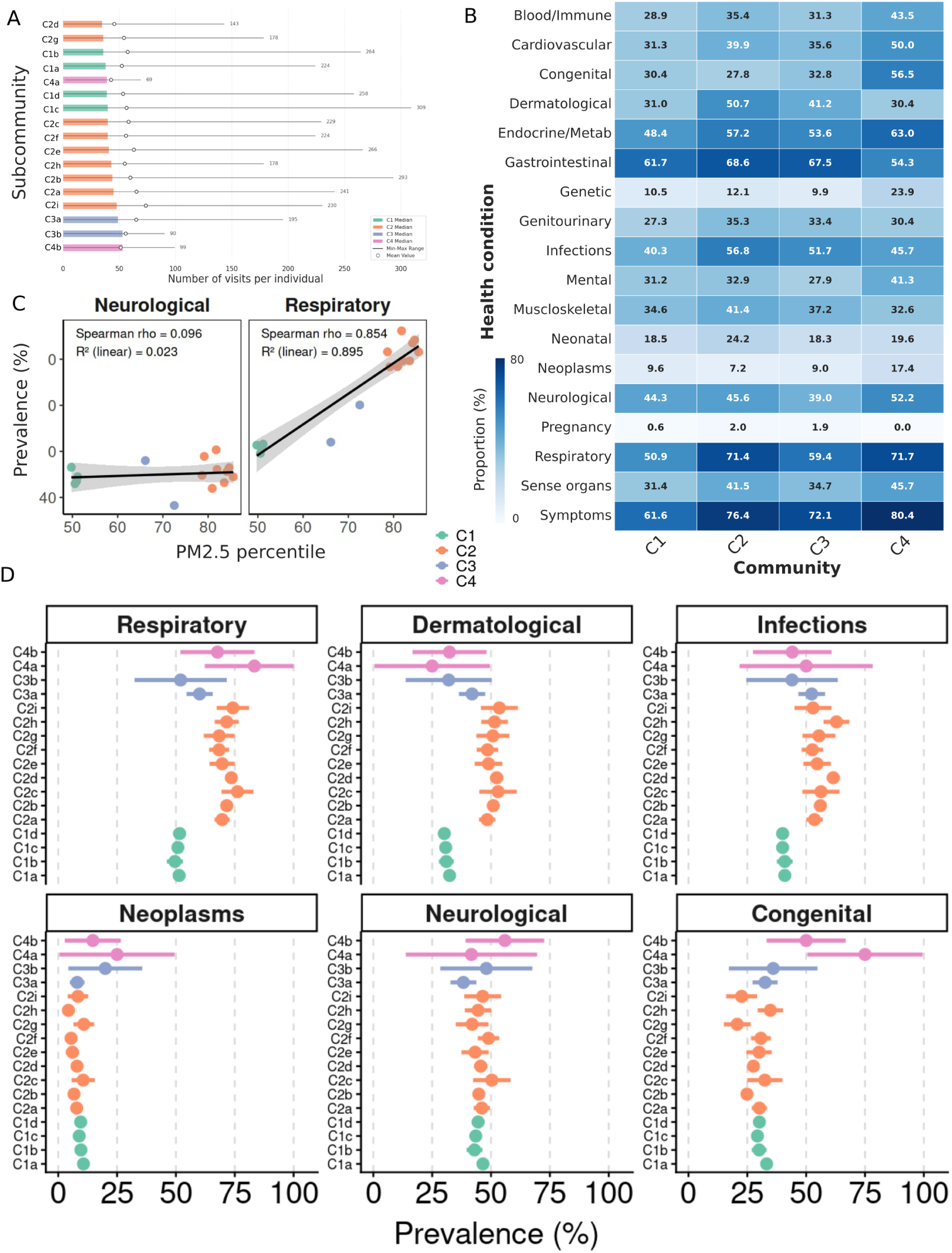
Health conditions across subcommunities. **(A)** Bar plot illustrating the median number of hospital visits per year in subcommunities. Mean and maximum are also indicated. **(B)** Heatmap showing the proportion of phenotypes coded by PheCodes at the main community level (C1, C2, C3, C4), with each cell reflecting the percentage of individuals displaying a specific category (e.g., Respiratory, Neurological, Infections, etc.). **(C)**Regression between prevalence and PM_2.5_ for Neurological and Respiratory conditions. Each dot represents one community. **(D)** Proportion of phenotypes at the subcommunity level and 95% confidence intervals, colored by community C1, C2, C3. Subcommunities from C4 were excluded due to the high uncertainty associated with fewer individuals (complete results are shown in Supplementary Figure S5).

In this Memphis cohort, C2 showed a particularly high prevalence of respiratory diseases (71.4%), infections (56.8%), and dermatological conditions (50.7%). In contrast, conditions with stronger genetic components (Huang et al., 2014; Lee et al., 2021), including neoplasms, neurological disorders, and congenital disorders, show more homogeneous distributions across communities. We evaluated disease prevalence across the aforementioned six categories at the subcommunity level. Although these categories are broad and warrant detailed case-by-case analysis, we observed a clear pattern: Communities showed variable prevalence for environmentally-driven diseases but consistent prevalence for genetically-driven diseases, suggesting that differential environmental exposures influence disease risk (**Figure 4D**). In line with what has been described so far, C2 has the largest variance across subcommunities in many cases, especially in congenital disorders that appeared generally similar across communities, with some cases, C2g and C2i, exhibiting a lower prevalence of congenital conditions.

Combining the population structure, environmental factors, and health conditions, we detected that fine particulate matter exposure was strongly associated with the respiratory disease burden across subcommunities: PM_2.5_ explained most of the variation in respiratory prevalence (Spearman *ρ* = 0.85, *R*^2^ = 0.90), whereas no meaningful association was observed for neurological disorders (*ρ* = 0.10, *R*^2^ = 0.02; **Figure 4C**).

Importantly, individuals across all subcommunities showed consistent healthcare access, with a median of 8.7 visits annually and 38 total visits per individual, suggesting that differences in disease patterns between communities are unlikely to be explained by differential access to care or under-diagnosis (**Figure 4A**).

### 2.5 Decomposing the sources of subcommunity health disparities

Up to this point, we have established that IBD-defined communities reflect ancestry and geography, with subcommunities sharing environmental exposures and showing enrichment for specific health conditions. Now, we ask whether we can take a further step by disentangling the sources of these differences. Specifically: *how much is attributable to measured environmental and socioeconomic factors, and how much remains unexplained?* To answer this, we implemented an interventional mediation analysis treating IBD subcommunity membership as the exposure, with age and sex as baseline covariates (**Figure 5A**). We focused on four pediatric-relevant outcomes selected *a priori* based on epidemiological evidence linking environmental exposures (PM_2.5_, poverty) to disease risk: asthma (n=1,757), dermatitis (n=742), epilepsy (n=1,502), and influenza (n=3,378) (**Figure 5B**). This focused, hypothesis-driven design prioritizes statistical precision in the mediation decomposition over exploratory breadth. For each outcome, we specified *a priori* a parsimonious set of neighborhood-level environmental mediators: PM_2.5_ air pollution and poverty (PPOV) rate for respiratory and dermatological conditions; obesity prevalence, smoking prevalence, and PPOV for infections.

**Figure 5:**
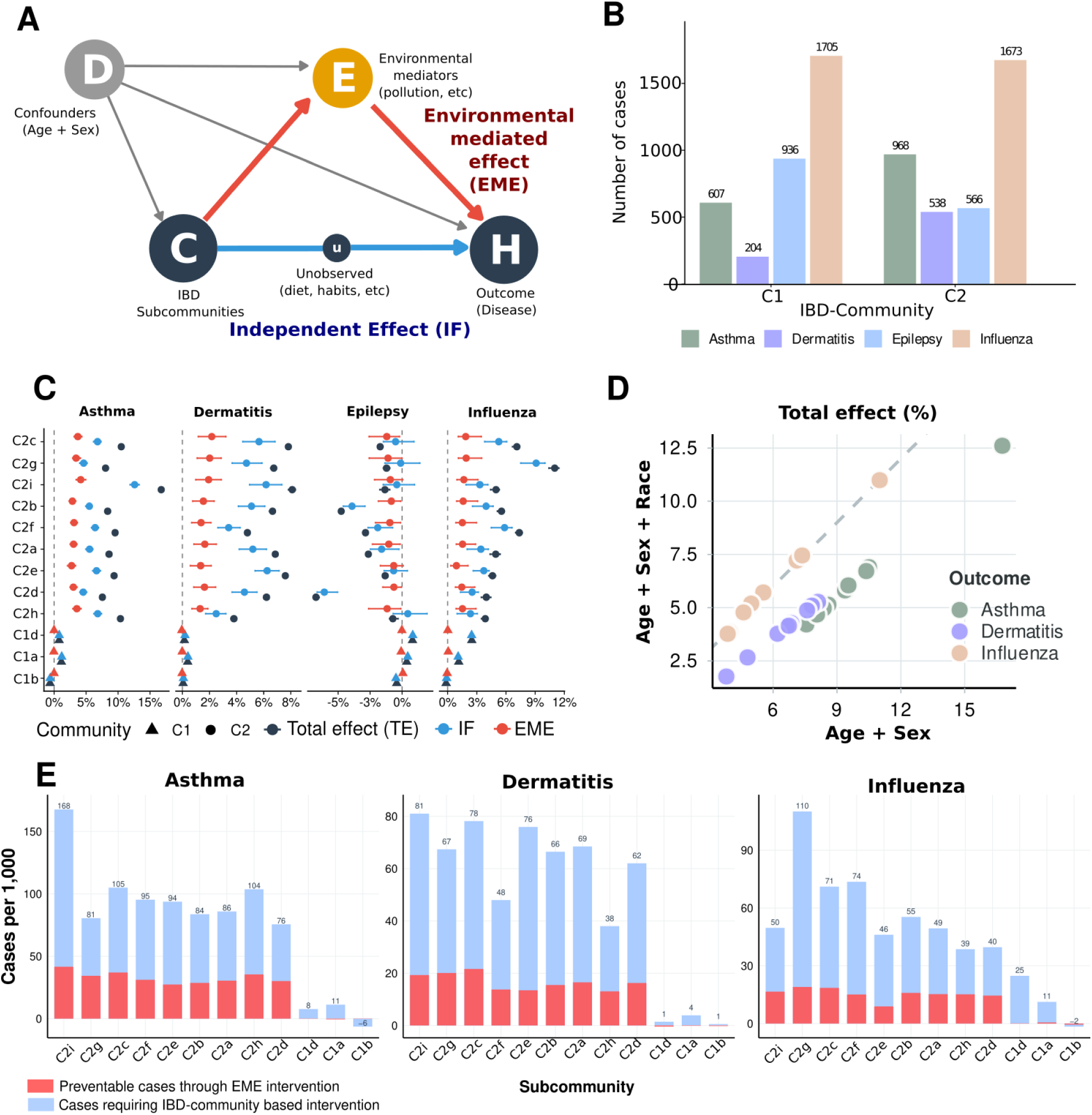
Prevalence of health conditions and decomposition of community effects on health. (A) Number of individuals with influenza, asthma, epilepsy, and dermatitis by community (C1–C3). (B) Directed acyclic graph (DAG) summarizing assumed relationships among baseline confounders (*D*), exposure (IBD subcommunity, *C*), mediators (environment, *E*), and outcome (health, *H*). (C) Risk difference (RD) relative to a reference community (*C*1*c*) for each condition, decomposed into the independent factor (IF; blue), the environmental mediated effect (EME; orange), and the total effect (TE; black); points denote estimates and horizontal bars indicate 95% confidence intervals. The vertical dashed line marks RD = 0. (D)Total effect for each subcommunity comparing the standard set of confounders (Age + Sex) with the addition of self-reported race. (E) Number to be treated analysis showing the number of individuals whose disease could be prevented through equalization of environmental exposures (EME), and the number requiring further investigation of IBD-community-specific factors (IF).

First, we estimated for each subcommunity the absolute difference of disease risk (risk difference, RD, or total effect) relative to subcommunity C1c, arbitrarily chosen as a reference, for comparative purposes. All nine C2 subcommunities exhibited a significantly elevated risk of disease prevalence for asthma (mean RD = +9.9 percentage points [pp]; range: 7.6–16.8 pp), dermatitis (mean RD = +6.5 pp; range: 3.8–8.1 pp) and influenza (mean RD = +5.9 pp; range: 3.9–11.0 pp), with 95% confidence intervals excluding zero (**Figure 5C**). Epilepsy showed an opposite pattern, with some C2 subcommunities exhibiting modestly lower risk near zero (mean RD = −2.6 pp; range: −6.7 to −0.7 pp).

We then separated the total risk difference (Total Effect, TE) into (i) an environmental mediated effect (EME): the portion of risk attributable to environmental mediators, i.e., PM_2.5_, poverty, and other SDoH variables exposure, (*C* → *E* → *H*), and (ii) an independent effect (IF): the residual component not explained by those mediators, and potentially explained by unmeasured factors captured by the IBD-community (*C* → *u* → *H*). Our data show that EME, explains only approximately one-third of the excess disease burden compared to two-thirds explained by IF. EME ranged from 2.7 to 4.2 pp for asthma, 1.3 to 2.2 pp for dermatitis, and 0.9 to 1.9 pp for influenza. On average across C2 subcommunities, environmental mediation accounted for 33% of the total effect for asthma, 26% for dermatitis, and 28% for influenza. In contrast IF, the residual risk elevation not explained by measured mediators, was substantial: 4.5 to 12.6 pp for asthma, 2.5 to 6.2 pp for dermatitis, and 2.3 to 9.1 pp for influenza. Subcommunity C2i exhibited the largest total and residual effects for asthma (TE = 16.8 pp; EME = 12.6 pp; IF = 4.2 pp), while C2g showed the highest effects for influenza (TE = 11.0 pp; EME = 9.1 pp; IF = 1.9 pp). On average, IBD-defined subcommunities retained more than two-thirds independent component (ranging from 67% for asthma to 74% for dermatitis) **(Supplementary Figure S7**). Overall, we concluded that the IF likely reflects a combination of genetic factors, unmeasured environmental exposures (e.g., diet, indoor air quality, healthcare utilization patterns), and social/cultural determinants that co-segregate with fine-scale IBD-based relatedness.

E-value analysis (VanderWeele and Ding, 2017) indicated that our findings are robust against moderate unmeasured confounding, with E-values for total effects ranging from 1.8 to 6.2 across outcomes and subcommunities. We additionally conducted a sensitivity analysis adjusting for self-reported race (**Figure 5D**, **Supplementary Figure S7**). Conditioning on self-reported race reduced statistical power; yet, several subcommunities retained significant direct and indirect effects (race adjustment attenuated total effects for asthma and dermatitis, but not for influenza). Because self-reported race may be partly downstream of community structure rather than an independent confounder, attenuation under race adjustment should be interpreted as bounding rather than identifying the share of variation captured by social labels. Overall, residual variation after adjusting for race, age, sex, and measured environmental factors suggests that IBD-defined community membership co-varies with unmeasured exposures relevant to clinical outcomes.

Finally, to gain insight from a public health perspective, we used the model (**Figure 5A**) to simulate interventions. If environmental exposures could be equalized across communities, approximately 33 asthma cases, 17 dermatitis cases, and 15 influenza cases could be prevented per 1,000 individuals in high-risk C2 subcommunities, which are concrete and actionable targets for intervention, as we show in (**Figure 5E**). However, these modifiable factors account for only part of the excess risk. The remainder, indexed by IBD-defined community structure, represents the maximum that could plausibly be attributed to community-specific factors, not a measurement of them. Because environmental exposures were observed at the ZIP-code level rather than per individual, environmental variation within neighborhoods is absorbed into this residual; with individual-level exposure data, part of it would be reassigned to the environment-mediated component.

### 2.6 Public health applications

To translate these findings into practice, we developed an open-source dashboard using BIG biobank data (https://github.com/MarsicoFL/BIG.Communities), which integrates community-specific prevalences, environment-mediated components, and the residual structure described above. For public health, this combined view reveals genetically related individuals across ZIP codes who share disease risk, enabling health departments to trace exposure pathways and quantify environmental versus unexplained contributions before intervening.

Finally, this network-based approach accommodates dynamic biobank growth, as demonstrated by our analysis incorporating 4,091 individuals from the HGDP-1kGP joint release (Koenig et al., 2024). It showed that existing community assignments remain stable upon the addition of new participants, and new communities arise capturing new genetic diversity (**Supplementary Figure S8**). Our application can be adapted to other cohorts, enabling joint exploration of IBD-based communities, health conditions, and environmental factors.

Similar approaches have demonstrated utility in other biobank initiatives, as described by Caggiano et al. (2023) and Dai et al. (2020).

## 3 Discussion

In this work, we have shown that IBD can be used to group individuals without relying on reference populations, and that in our cohort, the grouping is informative about environmental and population health factors. Our findings demonstrate that IBD-based clustering captures population structure at multiple scales, from continental ancestry patterns to fine-scale genetic subcommunities that reflect shared environmental exposures and social determinants of health. Beyond broad inter-community differences, we observed distinct health condition prevalence among subcommunities, suggesting that genetic relatedness correlates with both biological and environmental factors influencing disease risk. Importantly, incorporating IBD information into genotype-phenotype models accounts for both genetic similarity and shared environmental effects, providing a more comprehensive framework for association studies. Together, these results demonstrate the utility of considering both close and distant relationships between individuals for targeted public health interventions.

Recent discussions have focused on the risks of conflating race and ethnicity with genetic ancestry (Kozlov, 2024; National Academies of Sciences, Engineering, and Medicine, 2023; Popejoy, 2021; Carlson et al., 2022; Lewis et al., 2022, 2024). Typically, genetic studies categorize individuals based on continental-level labels; yet such classifications often overlook actual genetic relatedness. Our previous work demonstrated instances where individuals from different self-reported racial categories were closely related (i.e., second degree) (Buonaiuto et al., 2025). Similarly, ancestry-based classifications can separate genetically related individuals into distinct groups, particularly in cases involving admixed ancestry (Mas-Sandoval et al., 2023). In contrast, IBD-based hierarchical community detection allows for grouping individuals by their direct genetic relationships without relying on externally imposed labels based on arbitrarily defined reference populations. Although self-reported race is frequently used as the main proxy for unmeasured social factors (Manski et al., 2023), we show that residual variation in IBD-defined community structure, after adjusting for environmental and demographic confounders and self-reported race, reflects shared social and environmental exposures co-segregating with recent genealogy. Furthermore, unlike race and ethnicity, whose definitions vary across countries and lack universal applicability, IBD-based classification relies solely on genomic data, making it transferable across populations regardless of local demographic terminology. We emphasize that IBD-defined communities are clusters of recent genealogical relatedness, not racial or ethnic groups, even when they correlate with these descriptors due to historical demographic processes.

Recently, Shemirani et al. (2026) introduced continuous spectral components from IBD graphs for fine-scale population-structure adjustment in UK Biobank association studies, Belbin et al. (2021) and Caggiano et al. (2023) characterized IBD-based clusters in adult biobanks through broad phenome-wide scans, and Dai et al. (2020) demonstrated hierarchical IBD structure across the U.S. Genographic Project. Compared to the mentioned references, our framework addresses complementary aspects: (1) We focus on a pediatric biobank with neighborhood-level geocoding, (2) we build an explicit DAG-based decomposition (Figure 5A) of risk differences into known environmental factors-mediated and community-specific components under stated modeling assumptions, and (3) we provide a quantitative integration of social determinants of health (PM_2.5_, poverty, social vulnerability).

This work focuses on the entire cohort characterization; most of the analyses were based on the 17 main subcommunities that represent the predominant demographic structure. We show how subcommunity-level patterns emerge, providing information for risk assessment and public health interventions, and identifying groups that may be at risk for genetic and environmental factors. Furthermore, the network-based approach offers additional clinical utility by identifying and characterizing outlier individuals within minor subcommunities, allowing targeted investigation of health conditions potentially linked to their distinct fine-scale IBD-defined relatedness pattern (Caggiano et al., 2023; Lancaster et al., 2024).

In addition, population stratification, admixture, and shared environmental factors rep- resent significant sources of confounding in genome-wide association studies (GWAS) and admixture mapping, potentially leading to false positives and negatives (Tan and Atkinson, 2023; Devlin and Roeder, 1999; Caliebe et al., 2022; Liu et al., 2013). Addressing these confounders through methods such as principal component adjustment, mixed-model association analyses, and inclusion of local ancestry covariates has become critical to accurately identify true genotype-phenotype associations (Liu et al., 2013; Tan and Atkinson, 2023). Usually, in GWAS studies, only close relationships or ancestry are taken into account (Tan and Atkinson, 2023), and population structure adjustments are introduced by performing principal component analysis and incorporating the top principal components (PCs) as covariates. However, it has been observed that the introduction of PCs can lead to spurious associations in GWAS studies when including admixed populations (Grinde et al., 2024). The IBD-based grouping strategy inherently integrates distant genetic relatedness, ancestry, and shared environmental factors. Using IBD grouping to adjust population structure in association studies can be a promising line of future research.

A further consideration concerns the population descriptors conventionally used in genomic research. Self-reported race and ethnicity reflect the structure of U.S. census categories and EHR-coding practices, which do not capture the full continuum of self-identification (National Academies of Sciences, Engineering, and Medicine, 2023; Lewis et al., 2024). Inferred genetic ancestry, in turn, depends on reference panels such as HGDP-1kGP (Koenig et al., 2024), which under-represent Indigenous American, Middle Eastern, South Asian, and Oceanic diversity. These limitations motivate the IBD-based framework adopted here, which groups individuals based on their direct genealogical relationships, without imposing labels, reference panels, or pre-established social categories. Within this framework, genetic structure also carries a social character, possibly reflecting that kinship and shared social environments co-shape one another.

Despite its advantages, the IBD-based grouping approach has limitations. Its applicability may depend on the population density, the degree of local genetic structure, and the availability of genomic and geographic data (Tang et al., 2022). Future work should evaluate the integration of IBD-based categorization with existing correction strategies in GWAS (Zhou and Stephens, 2012) and gene-based association studies (Berrandou et al., 2023), particularly in cohorts with varying levels of endogamy, admixture, and environmental heterogeneity (Caliebe et al., 2022). Another aspect is the definition of relatedness, which lacks a universally accepted formulation. As discussed by Speed and Balding (2015), different measures, based on pedigree, allele, and segment sharing, reflect different interpretations of what it means to be related. In this study, we define relatedness as the total length of genome-shared IBD, a direct, quantitative measure of recent co-ancestry. Although often categorized for interpretability, this metric captures a continuous spectrum of genetic similarity.

Ultimately, our findings highlight how a deeper understanding of genetically defined communities can enhance the study of health disparities, uncover hidden confounders in genomic analyses, and open new avenues for integrating genetic and environmental factors into population-scale research.

## 4 Methods

### 4.1 Ethics

Our research followed the ethical guidelines established in the Declaration of Helsinki for human subjects’ medical research. We conducted this study according to approved ethical standards, with authorization from the University of Tennessee Health Science Center’s Institutional Review Board (IRB number: 23-09204-NHSR). All participants provided written informed consent, with parents or legal guardians giving consent for the children involved in the study. To protect privacy, we removed all identifying information from the data before conducting our analysis. Aggregated results, including community-level disease and exposure summaries, are made publicly available through the dashboard described above; individual-level community assignments are not returned to participants.

### 4.2 Sample Collection

Our study collected data from four key locations across Tennessee: (i) Le Bonheur Children’s Hospital (Memphis) serves a pediatric patient population that is predominantly self-identified as Black/African American. The hospital is located in a Memphis area shaped by historical residential segregation and persistent socioeconomic inequities. Recruitment began in October 2015 across all hospital settings. Patient distribution follows a distance gradient ((Buonaiuto et al., 2025)). Genomic DNA comes from leftover clinical blood linked to de-identified EHR data, with collection-consent discrepancies due to sample availability. (ii) Regional One Health (Memphis) joined in May 2022, focusing on adult genomic research from similar populations, with identical DNA collection procedures complementing LBCH’s pediatric focus. (iii) East Tennessee State University (Johnson City) added in May 2023 represents Appalachian populations through dedicated adult blood draws, supporting BIG’s commitment to rural communities. (iv) Family Resilience Initiative (Memphis), launched in January 2019, studies adverse childhood experiences through mother-child dyads, collecting biological samples quarterly over 18 months for DNA isolation, cortisol measurements, and clinical assessments to link biological and environmental data.

### 4.3 Clinical Data

BIG participants’ clinical information is extracted from Electronic Health Records (EHR) as flat files and transferred to UTHSC using secure protocols. This information encompasses demographics, hospital visits, diagnoses, procedures, medication details (both prescribed and administered), laboratory results, and vital measurements. We convert these elements into a limited data set (LDS) and align them with the OMOP (Observational Medical Outcomes Partnership) common data model. To facilitate analysis, we map ICD9/10 diagnosis codes to their corresponding PheCodes (Bastarache, 2021). Each PheCode was also associated to a broader category (Respiratory, Dermatological, Neurological, among others, see Supplementary Figure S5). Disease conditions were defined using specific PheCodes: asthma was identified using PheCode RE_475; dermatitis using PheCodes beginning with 690, which includes various subtypes of skin inflammation conditions.

### 4.4 DNA Sequencing, quality control and variant analysis

All samples were sequenced on an Illumina NovaSeq 6000 platform. The samples were processed with NEB/Kapa reagents, captured with the Twist Comprehensive Exome Capture design, enhanced by Regeneron-designed spikes targeting sequencing genotyping sites. Among the sequenced samples, 95.2% achieved an average sequencing depth of at least 20X, and 99.3% of the samples had more than 90% of their bases covered at 20X or greater, highlighting the overall quality of the data. The genotyping spike targets an additional ≈1.4M variants in the human genome. Genotyping call rate (percentage of SNP / indels targeted genotyping at which a call can be made) is 99.0%. All samples were sequenced on an Illumina NovaSeq 6000 system on S4 flow cells sequencer using 2×75 paired-end sequencing.

Sequence reads were aligned by the Burrows-Wheeler Aligner (BWA) MEM (Li and Durbin, 2009) to the GRCh38 assembly of the human reference genome in an alt-aware manner. Duplicates were marked using Picard, and mapped reads were sorted using sambamba (Tarasov et al., 2015). DeepVariant v0.10.0 with a custom exome model was used for variant calling (Poplin et al., 2018), and the GLnexus v1.2.6 tool was used for joint variant calling (Lin et al., 2018). The variants were annotated using a Variant Effect Predictor (VEP 110) (McLaren et al., 2016). Phasing was performed using ShapeIT v5 (Delaneau et al., 2013). Our dataset comprised 6,886,631 variable sites after quality control, combining both exome capture and targeted sequencing data.

### 4.5 Ancestry Inference

To characterize genetic ancestry in the BIG cohort, we inferred global ancestry with RFMix (Maples et al., 2013) using methodology described in (Buonaiuto et al., 2025). Reference samples from 1000 Genomes and HGDP (Koenig et al., 2024) were merged with BIG participants, and quality-controlled references (Mas-Sandoval et al., 2023) were filtered using ADMIXTURE (k = 4) (Alexander and Lange, 2011) to retain samples with major ancestry proportion *>*0.99, defining African (AFR), American (AMR), European (EUR), and East Asian (EAS) ancestries. Four-way deconvolution used RFMix with standard parameters (5 terminal nodes, 12 generations, 10 EM iterations); CSA ancestry was excluded due to negligible proportions. Discrete categories were defined as: *>*85% single ancestry for unadmixed groups; *>*15% from two ancestries summing to *>*85% for two-way admixed (EUR-AMR, EUR-AFR); and *>*15% from three or more ancestries for Multiway, with the 85% threshold reflecting that contributions *>*15% are accurate while lower values show higher error (Gravel, 2012; Sohail et al., 2023).

### 4.6 Identity-by-descent

To identify identity-by-descent (IBD) segments, we used hap-ibd in the phased dataset comprising 13,143 genomes from the BIG cohort and 4,091 genomes from the HGDP-1kGP, focusing exclusively on autosomal loci (Zhou et al., 2020). Hap-ibd was executed with a minimum seed parameter of 2 cM to focus on the detection of IBD segments reflecting recent common ancestry, given that segments of this length or greater provide information for fine-scale relationship inference (Huff et al., 2011).

Post-processing protocols established by Browning and Browning (2013; 2015) were applied to refine the detected segments (using Refined IBD (Browning and Browning, 2013)). Specifically, the merge-ibd-segments tool was employed with default parameters, consolidating fragmented segments to improve continuity. Additionally, gaps containing at most one discordant homozygote and segments shorter than 0.6 cM were removed, significantly enhancing detection accuracy and efficiency in large-scale genomic datasets. The 2 cM seed threshold was retained to maximize detection of recent shared ancestry; classical false-positive rates reported for short segments (Henn et al., 2022) are reduced by the post-processing protocols applied above (gap-merging, low-quality segment filtering, and exclusion of high-LD regions).

For quality control, segments overlapping centromeres, telomeres, the human leukocyte antigen region, and low complexity regions were excluded. To identify genome regions with a high shared genetic material, we calculated total identity by descent at each marker position and removed those that deviated significantly from the genome-wide average. For downstream analysis, IBD segment lengths were summed across all chromosomes between each pair of individuals, generating the total shared IBD. Finally, only for illustrative purposes, we classified the degree of relationships based on IBD sharing (Huff et al., 2011; Gusev et al., 2012; Browning and Browning, 2015) using the pedsuite R package (Vigeland, 2021).

### 4.7 Network, community detection and topology

A genetic network was constructed in which each BIG participant was represented as a node, and edges were defined based on the total genome-wide IBD segment sharing. Edge weights were proportional to the total IBD shared between individuals. To build this network, pairwise IBD data were processed and filtered, excluding close-kin pairs following Dai et al. (2020). For visualization and exploratory analyses, the network was represented using the Fruchterman-Reingold layout.

We evaluated the Louvain and Leiden algorithms for community detection (Anuar et al., 2021). Both were implemented using a multilevel approach. This means that the algorithms were run iteratively, initially identifying broad-scale genetic structures and subsequently resolving increasingly finer-scale subcommunities. Comparative analysis revealed an Adjusted Rand Index (ARI) of 0.975, indicating high similarity between community assignments from both methods. However, we selected the Leiden algorithm due to its improved resolution, robustness, and ability to avoid disconnected communities, a known limitation of Louvain (Anuar et al., 2021).

The hierarchical Leiden algorithm was applied, with modularity as the objective function, using a resolution parameter of 0.3 and a beta value of 2. These values were determined empirically to optimize the balance between identifying meaningful broad-scale community structures and maintaining balanced sizes of subcommunities. Sensitivity analyses varying the resolution parameter (0.1–0.5) and beta values (1–3) yielded qualitatively similar community structures (ARI > 0.92 across configurations), with the reported parameters. This approach aligns conceptually with previously published methods for hierarchical community detection based on IBD networks (Dai et al., 2020), but it explicitly specifies parameter values adapted to our dataset characteristics and study objectives.

Genetic differentiation between communities was independently assessed using the fixation index (F_ST_), calculated with PLINK. This analysis quantitatively characterized genetic distances between communities, offering independent validation and additional genetic context for the hierarchical community structures identified by the Leiden algorithm.

Network topology analysis was performed using NetworkX to characterize the structural properties of IBD-based genetic networks within each community partition. For each individual node, the connectivity ratio was computed as the normalized degree centrality, calculated by dividing the number of direct connections by the mean degree within the respective community. Betweenness was quantified using the standard algorithm that measures the fraction of shortest paths between all node pairs that traverse through each individual node, subsequently normalized by the community-specific mean betweenness score to facilitate comparative analysis across communities with different topological characteristics. The normalization procedure ensures that values above 1.0 indicate above-average connectivity or centrality within the community context, while values below 1.0 represent below-average network importance. To be consistent and informative in our context, network topology analyses require more than 30 nodes; for this reason, C4 subcommunities were not included.

To contextualize the genetic structure of the BIG cohort and validate the robustness of the method to the inclusion of new participants, we performed a sensitivity analysis incorporating 4,091 genomic sequences from the joint HGDP-1000GP resource (Koenig et al., 2024). In this combined dataset (n=17,234 individuals), pairwise IBD was recalculated and processed using identical parameters and filtering thresholds as the BIG-only analysis. The same hierarchical Leiden algorithm was applied S8.

The first iteration identified ten communities, four of which (C1–C4) contained the vast majority of BIG individuals, corresponding to the previously applied BIG-only approach. Subsequent iterations revealed 72 subcommunities in total, with C2 exhibiting the highest number of subcommunities, followed by C4, C1, and C3. At the subcommunity level, classification remained concordant with the BIG-only analysis. Notably, many subcommunities were composed specifically of HGDP-1000GP individuals, demonstrating the model’s sensitivity to detecting novel communities of individuals genetically distant from those currently identified in BIG. The number of subcommunities did not correlate directly with parent community size, suggesting the influence of genetic diversity rather than sample size alone.

### 4.8 Geographical analysis

To analyze the geographical patterns of genetic relationships, we geolocated BIG cohort participants using ZIP code at neighborhood level data. Distances between neighborhoods were calculated using the sf R package (Pebesma and Bivand, 2018), and therefore, we obtained a matrix of distances between individuals (with zero as the distance between those who shared the same neighborhood). A network based on the geographical distance was constructed, representing the subcommunities in the nodes and the inverse of the median distance for the edges. Geographical data was integrated with the 24th USA census (2020)-based metrics. It provided several demographic measures and also social determinants of health.

### 4.9 Population descriptors

We use three orthogonal descriptors throughout this work, distinguished operationally and conceptually:

(i) *Self-reported race and ethnicity (SIRE)* are social and administrative categories extracted from electronic health records, used as proxies for shared social environment and exposure to structural inequities, not as biological variables (National Academies of Sciences, Engineering, and Medicine, 2023; Lewis et al., 2024). Hospital staff selected self-reported race categories from a dropdown menu following HL7 v2.5 standards (https://hl7-definition.caristix.com/v2/HL7v2.5/Tables/0005); the system allowed multiple self-reported race codes for individuals identifying with more than one background. Due to inconsistencies in legacy records, free-text and dropdown entries were harmonized to HL7 v2.5 categories while preserving self-identification (Supplementary Table S1). Hispanic/Latino ethnicity was recorded separately from race.
(ii) *Inferred genetic ancestry* refers to RFMix-derived ancestry proportions estimated against the harmonized HGDP-1kGP reference panel (Koenig et al., 2024), with components labeled AFR, AMR, EUR, and EAS. These labels denote reference-panel ancestry components and not biological or social identities; in particular, AMR refers to the admixture-derived component in the 1000G reference and should not be conflated with Indigenous American identity, which our reference panel cannot resolve.
(iii) *IBD-defined communities* are network-derived clusters of recent genealogical relatedness, obtained by hierarchical Leiden community detection on the IBD-sharing network. Communities (C1–C4) and subcommunities are clusters of pairwise relatedness, not racial, ethnic, or ancestry groups, even when they correlate with these descriptors due to historical demographic processes.

### 4.10 Social determinants of health

In our analysis, each subcommunity is characterized by aggregating Social Determinants of Health (SDoH) variables by neighborhood level. Therefore, we were able to assign SDoH to each BIG participant based on zip code. We analyzed SDoH variables from multiple sources. U.S. Census (2020) provided population density, poverty rates and Gini index. CDC contributed the Social Vulnerability Index (2020), obesity prevalence and smoking rates (2021). Geographic factors included USDA urban/rural designations (2019) and Treasury Department/CDFI metro classifications (2020). We incorporated unemployment rates (Treasury Department/CDFI, 2016-2020), and EPA’s PM_2.5_ particulate matter percentiles (2024).

We acknowledge that neighborhood-level assignment of exposures (an instance of ecological inference, with associated modifiable-areal-unit limitations) may not fully capture individual-level variation, particularly in cities with marked residential segregation. However, for pediatric populations with limited residential mobility, ZIP code-level metrics provide reasonable proxies for cumulative environmental exposure. Critically, ZIP-averaged measurement of the mediators introduces classical measurement error in the mediation path, which attenuates the environmentally-mediated component (EME) and correspondingly inflates the independent factor (IF) in the decomposition. The IF estimates reported here should therefore be interpreted as an upper bound on the genuinely community-specific contribution; refinement to individual-level exposure data would shift mass from IF toward EME.

Continuous variables (e.g., *PM*_2_._5_, *CSMOKING*) are averaged between individuals, while categorical variables (*A*9_*SV I*) are converted to ordinal scores by appropriate mapping and are similarly averaged. Each community is represented by a profile vector, and the Euclidean distance between communities *i* and *j* is calculated as

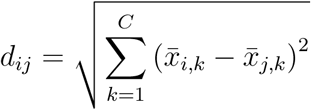

where *x̄_i,k_* is the mean value of the variable SDoH *k* for the community *i* and *x̄_j,k_* the mean value of same variable for the community *j*. The distance is normalized into a similarity metric by

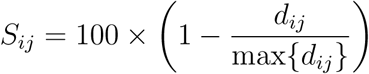

ensuring that identical community profiles yield a similarity of 100%.

### 4.11 Interventional effect analysis

We estimated interventional direct and indirect effects (Pearl, 2001; VanderWeele and Vansteelandt, 2014; Vansteelandt and Daniel, 2017) for IBD subcommunity (*C*; reference *C*1*c*) on binary health outcomes (*H*) using g-computation (Robins, 1986) on the risk-difference scale. Baseline covariates (*D*) included age and sex (plus self-reported race in sensitivity analyses). Environmental mediators (*E*) were selected *a priori* based on epidemiological evidence (Johnson et al., 2021; Mersha and Abebe, 2015) and assessed for collinearity prior to model specification: for asthma and dermatitis, *E* = {PM_2_._5_, PPOV}; for influenza, *E* = {OBESITY, CSMOKING, PPOV}; for epilepsy, *E* = {UNEMPLOY, OBESITY, PPOV}.

The interventional decomposition adopted here partitions risk differences under identification assumptions that we cannot test from these observational data. IBD-defined community membership (*C*) is a stable, genealogically determined characteristic that does not vary with outcomes, ruling out reverse association from *H* to *C*. Neighborhood exposures (*E*) at ZIP-code resolution were largely fixed during the period in which pediatric outcomes accumulated. Unmeasured factors, including individual mobility, indoor exposures, healthcare utilization, and cultural or behavioral characteristics, may confound the mediator-outcome relationship and are not addressed by our specification. We therefore interpret the resulting partition (TE, EME, IF) descriptively, as the decomposition of risk differences that holds under the specified DAG and identification assumptions; the E-value sensitivity analysis (VanderWeele and Ding, 2017) characterizes the strength of unmeasured confounding required to overturn the observed patterns.

Estimation used cross-fitted g-computation (*K* = 4 folds). In each fold we fit (i) a logistic outcome regression *q*(*C, D, E*) with natural splines for continuous covariates and factor indicators for categorical covariates, and (ii) a joint mediator model via sequential factorization (Imai et al., 2010) using Gaussian GLMs on automatically selected transformation scales (logit for bounded mediators, log for right-skewed, identity otherwise).

The g-formula requires integrating over the mediator distribution, which lacks a closed-form solution. We approximated these integrals via Monte Carlo simulation (500 draws), sampling mediator values by adding resampled residuals to predicted conditional means on transformed scales, then propagating through the outcome model. Counterfactual risks were averaged over simulations and individuals. Uncertainty was quantified via community-cluster weighted bootstrap (500 replicates), drawing *w_c_* ∼ Exp(1) per subcommunity and re-estimating effects with weighted likelihood to account for within-cluster correlation. We report percentile-based 95% confidence intervals. Analyses excluded subcommunities with *n <* 100 and C3a (positivity violations). Missing data were handled via rare-category lumping for categorical variables and median imputation with missingness indicators for continuous variables.

We conducted E-value sensitivity analysis (VanderWeele and Ding, 2017) to assess robustness to unmeasured confounding. E-values were computed by converting risk differences to approximate risk ratios under baseline prevalence estimates from the reference community, then applying the standard formula.

Additionally, we performed a specification sensitivity analysis adjusting for self-reported race, used here as a proxy for accumulated social exposures rather than as a biological covariate (National Academies of Sciences, Engineering, and Medicine, 2023; Lewis et al., 2024; Manski et al., 2023), to evaluate whether IBD-defined community effects persist beyond what self-reported race captures.

To estimate the public health impact of equalizing environmental exposures, we calculated the number of preventable cases per 1,000 individuals as the environmental mediated effect (EME) multiplied by 1,000. This represents the expected reduction in disease burden if neighborhood-level exposures (e.g., PM_2.5_, poverty) in high-risk subcommunities were equalized to reference levels.

### 4.12 Software availability

All analysis code, including IBD calling, network construction, community detection, and the interactive dashboard, is available at https://github.com/MarsicoFL/BIG.Communities and permanently archived on Zenodo (https://doi.org/10.5281/zenodo.19766414). The pipeline used the following software: BWA-MEM (Li and Durbin, 2009), Picard, sambamba (Tarasov et al., 2015), DeepVariant v0.10.0 (Poplin et al., 2018), GLnexus v1.2.6 (Lin et al., 2018), Variant Effect Predictor v110 (McLaren et al., 2016), ShapeIT v5 (Delaneau et al., 2013), hap-ibd (Zhou et al., 2020), RFMix (Maples et al., 2013), ADMIXTURE (Alexander and Lange, 2011), PLINK, the Leiden algorithm (Anuar et al., 2021), NetworkX, the sf R package (Pebesma and Bivand, 2018), and the pedsuite R package (Vigeland, 2021).

## 5 Data availability

Individual-level genomic and clinical records analyzed in this study are controlled-access under the BIG biorepository’s IRB protocol (University of Tennessee Health Science Center IRB 23-09204-NHSR), in compliance with HIPAA and institutional requirements protecting participants of a predominantly pediatric cohort. The following derived outputs supporting the findings of this paper are publicly available without restriction at https://github.com/MarsicoFL/BIG.Communities: aggregated ancestry proportions per community and subcommunity, de-identified IBD-network community assignments, neighborhood-level social-determinants-of-health summaries, and risk-difference estimates from the mediation analyses. The interactive dashboard built on these outputs is available at https://francomarsico.shinyapps.io/BIG_Communities/. All analysis code is permanently archived on Zenodo (https://doi.org/10.5281/zenodo.19766414). Researchers seeking access to individual-level data may submit a request to the BIG Research Oversight Committee at https://uthsc.edu/cbmi/big/ or; requests are reviewed within four weeks, and approved access is granted for the duration of the proposed project under a data use agreement.

## Regeneron Genetics Center

### RGC Management and Leadership Team

Aris Baras, Gonçalo Abecasis, Adolfo Ferrando, Giovanni Coppola, Andrew Deubler, Luca A Lotta, John D Overton, Jeffrey G Reid, Alan Shuldiner, Katherine Siminovitch, Jason Portnoy, Marcus B Jones, Lyndon Mitnaul, Alison Fenney, Jonathan Marchini, Manuel Allen Revez Ferreira, Maya Ghoussaini, Mona Nafde, William Salerno, Cristen Willer, Lourdes Crane.

### Sequencing and Lab Operations

John D Overton, Christina Beechert, Erin Fuller, Laura M Cremona, Eugene Kalyuskin, Hang Du, Caitlin Forsythe, Zhenhua Gu, Kristy Guevara, Michael Lattari, Alexander Lopez, Kia Manoochehri, Prathyusha Challa, Manasi Pradhan, Raymond Reynoso, Ricardo Schiavo, Maria Sotiropoulos Padilla, Chenggu Wang, Sarah E Wolf.

### Genome Informatics and Data Engineering

Jeffrey G Reid^5^, Mona Nafde^5^, Manan Goyal^5^, George Mitra^5^, Sanjay Sreeram^5^, Rouel Lanche^5^, Vrushali Mahajan^5^, Sai Lakshmi Vasireddy^5^, Gisu Eom^5^, Krishna Pawan Punuru^5^, Sujit Gokhale^5^, Shehroze Aamer^5^, Pooja Mule^5^, Mudasar Sarwar^5^, Muhammad Aqeel^5^, Xiaodong Bai^5^, Lance Zhang^5^, Sean O’Keeffe^5^, Razvan Panea^5^, Evan Edelstein^5^, Ayesha Rasool^5^, William Salerno^5^, Evan K Maxwell^5^, Boris Boutkov^5^, Alexander Gorovits^5^, Ju Guan^5^, Alicia Hawes^5^, Olga Krasheninina^5^, Samantha Zarate^5^, Adam J Mansfield^5^, Lukas Habegger^5^, Stephen Tahan^5^.

### Analytical Genetics and Data Science

Gonçalo Abecasis^5^, Manuel Allen Revez Ferreira^5^, Joshua Backman^5^, Kathy Burch^5^, Adrian Campos^5^, Liron Ganel^5^, Sheila Gaynor^5^, Benjamin Geraghty^5^, Arkopravo Ghosh^5^, Christopher Gillies^5^, Lauren Gurski^5^, Eric Jorgenson^5^, Tyler Joseph^5^, Michael Kessler^5^, Jack Kosmicki^5^, Adam Locke^5^, Priyanka Nakka^5^, Jonathan Marchini^5^, Karl Landheer^5^, Olivier Delaneau^5^, Maya Ghoussaini^5^, Anthony Marcketta^5^, Joelle Mbatchou^5^, Jonathan Ross^5^, Carlo Sidore^5^, Eli Stahl^5^, Timothy Thornton^5^, Rujin Wang^5^, Kuan-Han Wu^5^, Bin Ye^5^, Blair Zhang^5^, Andrey Ziyatdinov^5^, Yuxin Zou^5^, Jingning Zhang^5^, Kyoko Watanabe^5^, Mira Tang^5^, Frank Wendt^5^, Suganthi Balasubramanian^5^, Suying Bao^5^, Kathie Sun^5^, Chuanyi Zhang^5^, Sean Yu^5^, Aaron Zhang^5^, David Corrigan^5^, Dhruv Shidhaye^5^, Chen Wang^5^, Keyrun Adhikari^5^, Alexander Lachmann^5^, Anna Alkelai^5^, Mark Weiner^5^.

### Therapeutic Area Genetics

Adolfo Ferrando^5^, Giovanni Coppola^5^, Luca A. Lotta^5^, Alan Shuldiner^5^, Katherine Siminovitch^5^, Brian Hobbs^5^, Jon Silver^5^, William Palmer^5^, Rita Guerreiro^5^, Amit Joshi^5^, Antoine Baldassari^5^, Cristen Willer^5^, Sarah Graham^5^, Ernst Mayerhofer^5^, Erola Pairo Castineira^5^, Mary Haas^5^, Niek Verweij^5^, George Hindy^5^, Jonas Bovijn^5^, Tanima De^5^, Luanluan Sun^5^, Olukayode Sosina^5^, Arthur Gilly^5^, Peter Dornbos^5^, Moeen Riaz^5^, Manav Kapoor^5^, Gannie Tzoneva^5^, Veera Rajagopal^5^, Sahar Gelfman^5^, Vijay Kumar^5^, Jacqueline Otto^5^, Jose Bras^5^, Silvia Alvarez^5^, Jessie Brown^5^, Hossein Khiabanian^5^, Joana Revez^5^, Kimberly Skead^5^, Jae Soon Sul^5^, Lei Chen^5^, Sam Choi^5^, Amy Damask^5^, Nan Lin^5^, Charles Paulding^5^, Sameer Malhotra^5^, Joseph Herman^5^, Jacob McPadden^5^, David Blair^5^.

### Research Program Management and Strategic Initiatives

Marcus B Jones^5^, Michelle G LeBlanc^5^, Nadia Rana^5^, Jennifer Rico-Varela^5^, Jaimee Hernandez^5^, Larizbeth Romero^5^, Ashley Paynter^5^.

### Senior Partnerships and Business Operations

Randi Schwartz^5^, Lourdes Crane^5^, Alison Fenney^5^, Jody Hankins^5^, Anna Han^5^, Samuel Hart^5^, Ryan Smith^5^.

### Business Operations and Administrative Coordinators

Ann Perez-Beals^5^, Gina Solari^5^, Johannie Rivera-Picart^5^, Michelle Pagan^5^, Sunilbe Siceron^5^. ^5^Regeneron Genetics Center, Tarrytown, NY, USA.

---

**Figure S1:**
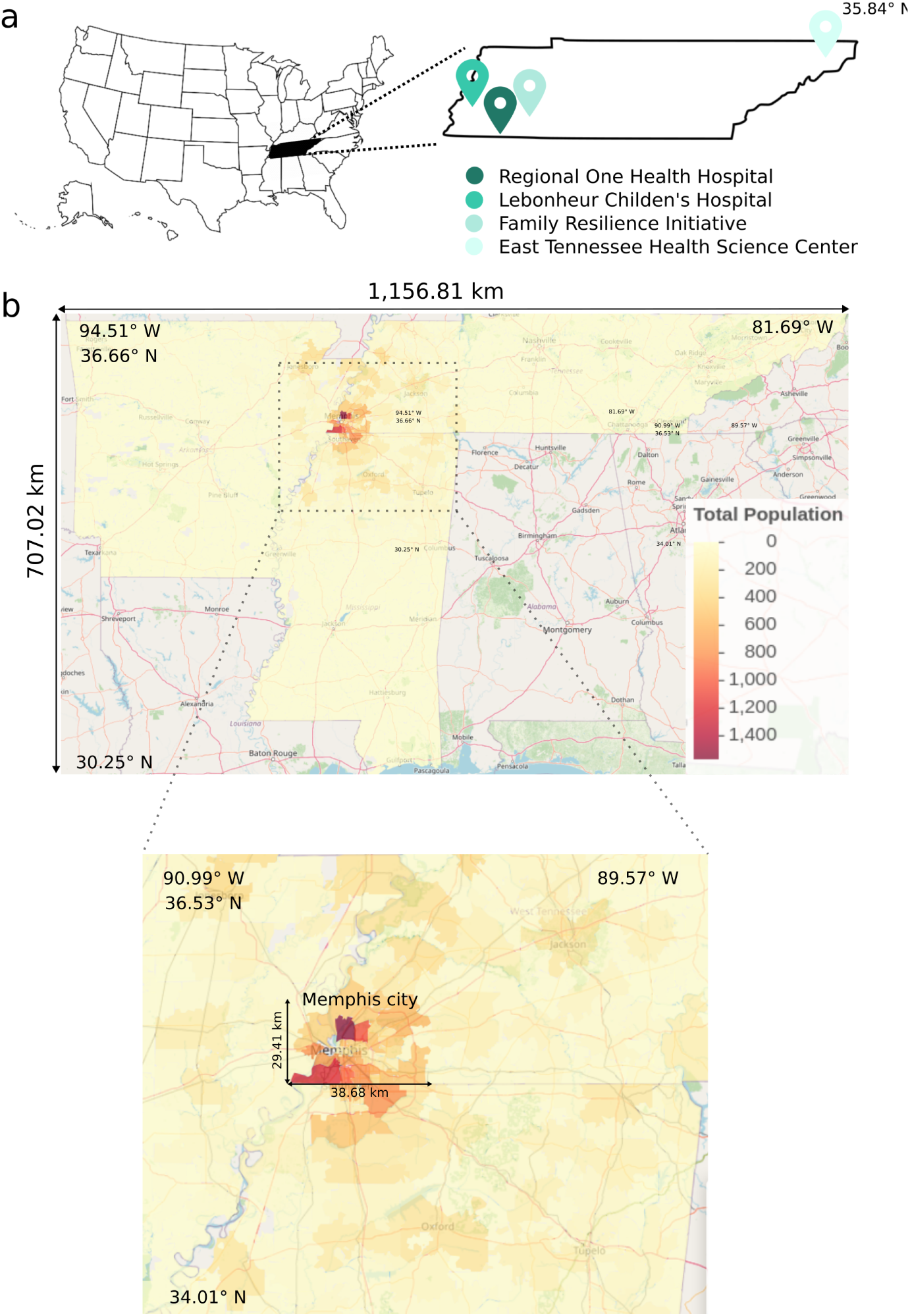
Demography of enrolled participants. (a) The four sample sites (b) Number of enrolled participants by ZIP code. The region surrounding Memphis City is zoomed in, and distances in Km are shown. Maps were produced with the leaflet package (v. 2.2.1) using GeoJSON data for state ZIP-code boundaries publicly available.

**Figure S2:**
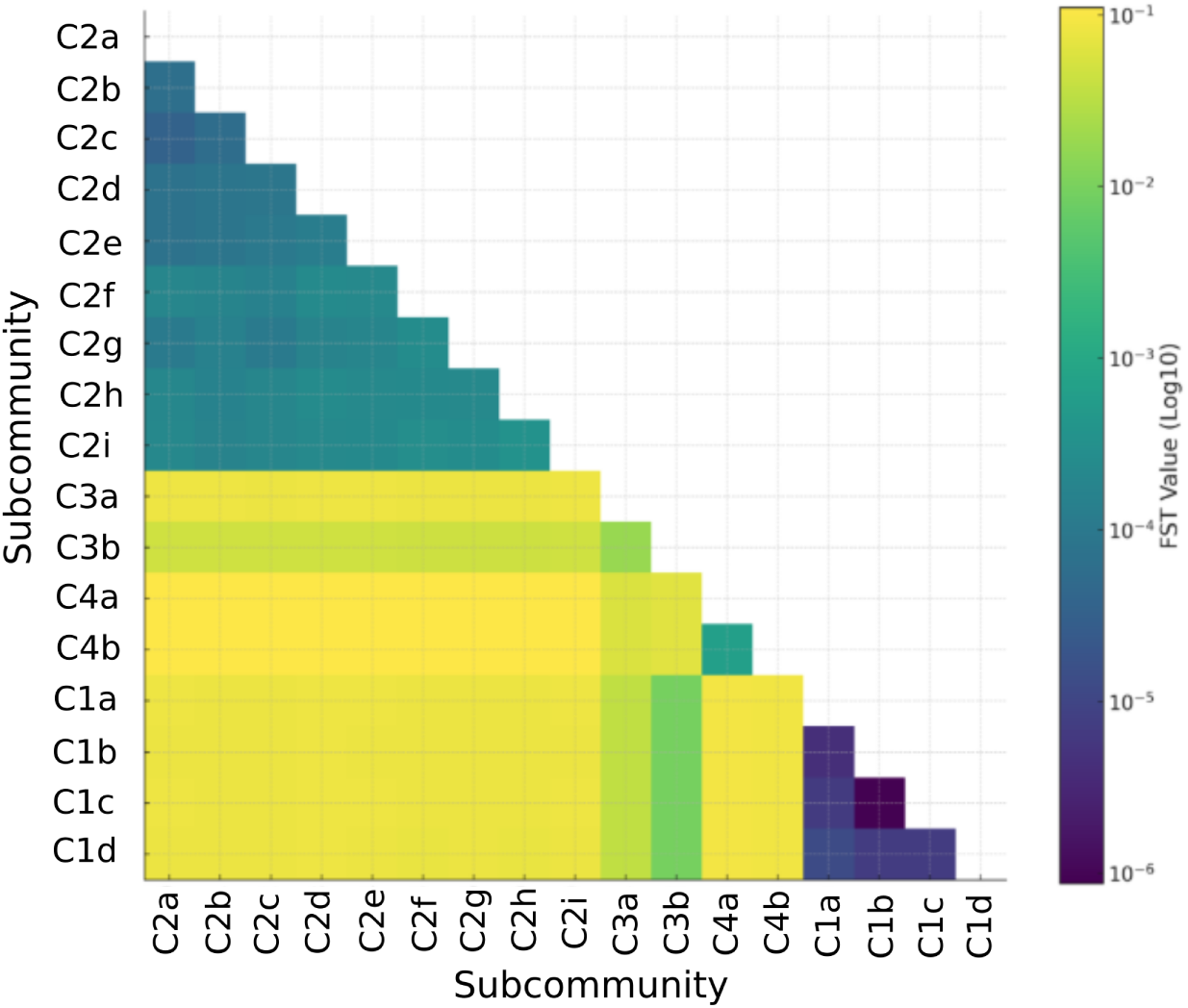
Pairwise Fst values between subcommunities. This analysis was performed using PLINK. It can be seen that subcommunities within C1 have the lowest fst values when analyzed each other. Contrastingly, C2 show higher values, indicative of more genetic diversity in the group.

**Figure S3:**
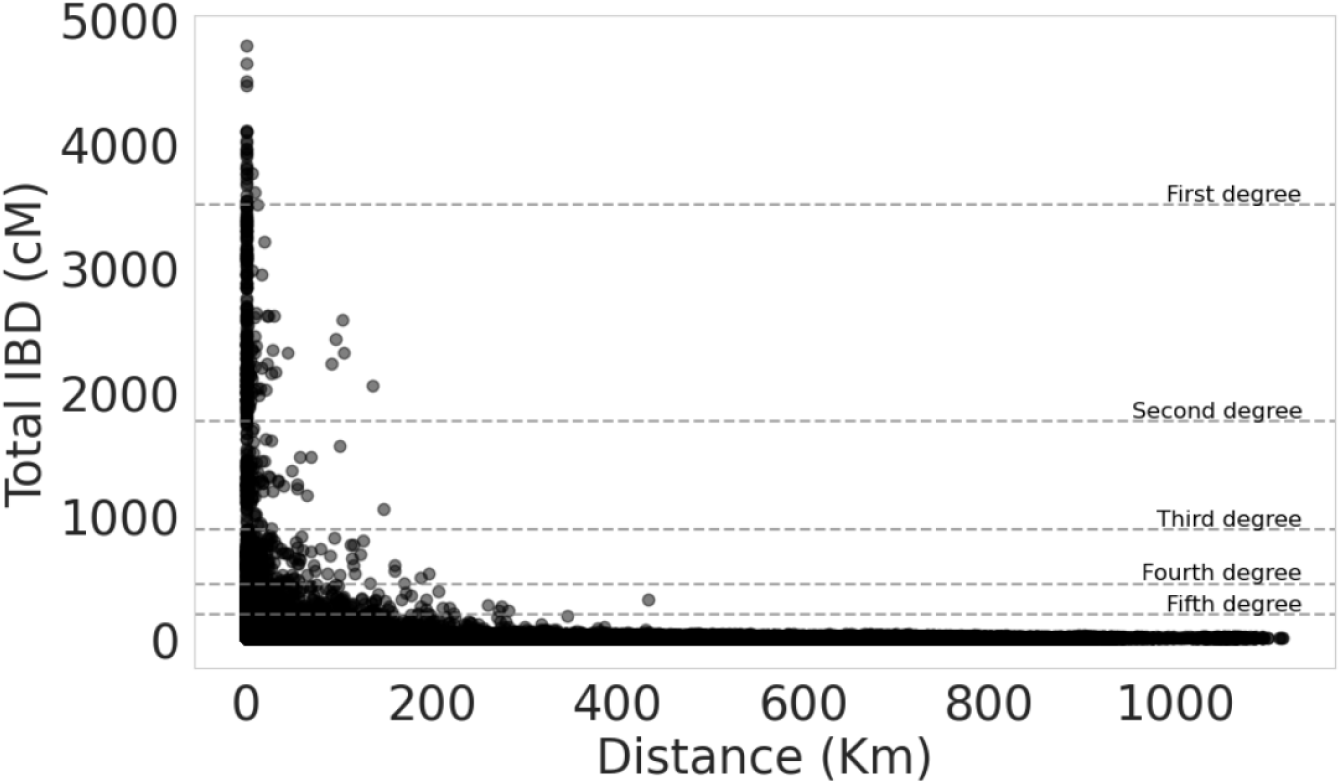
Total IBD (cM) and Distance (Km). Different cutoffs for degrees of relationships are shown. The plot elucidates how higher total IBD values are related with closer distances. Geographical distances have been computed with sf R package (Pebesma and Bivand, 2018).

**Figure S4:**
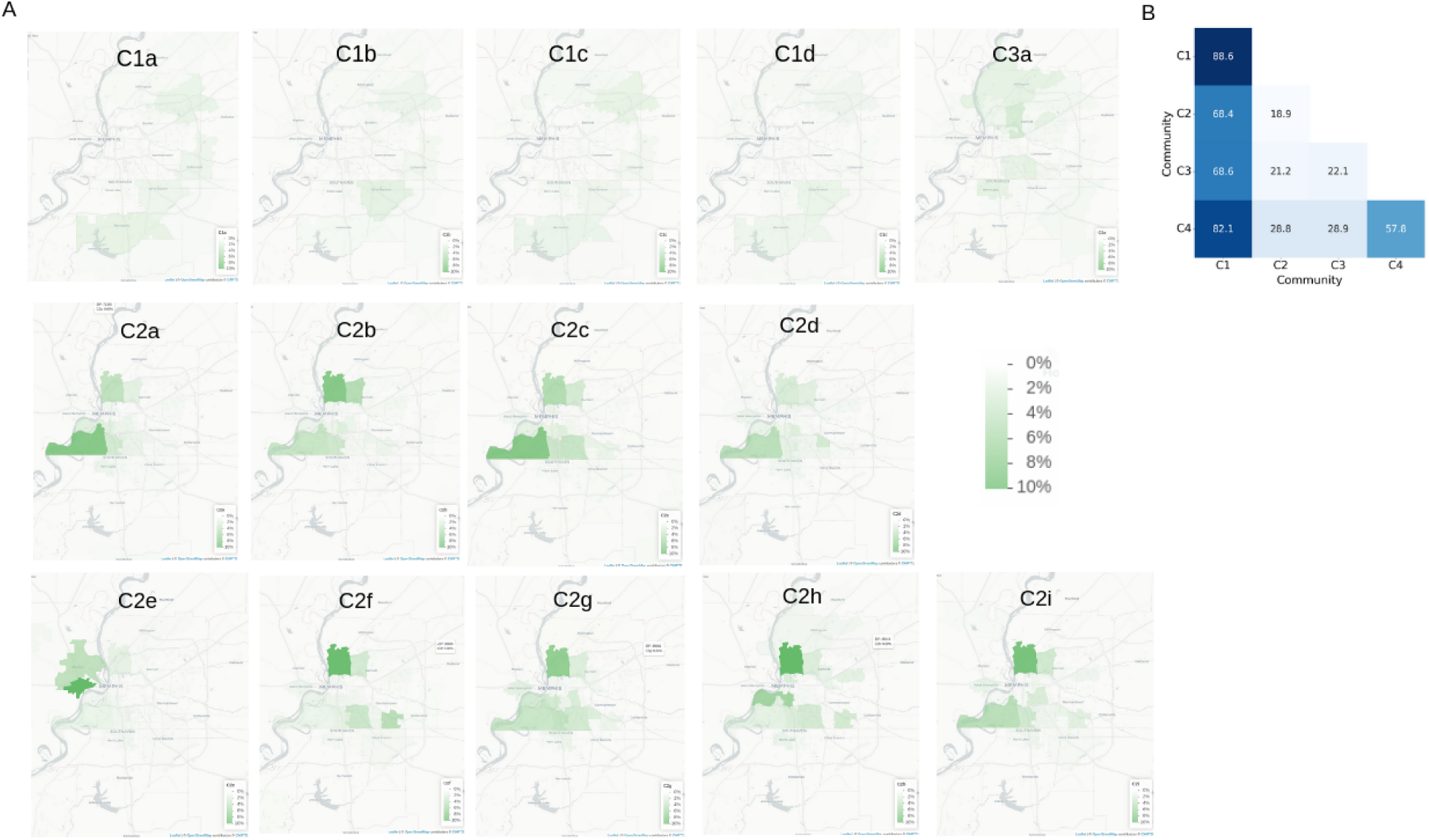
Geographical distribution of subcommunities around Memphis area. A. Maps centered in Memphis city (34.96°-35.36°N: 88.70°-90.70° W) showing the proportional distribution of individuals from each subcommunity across zip codes. Maps were produced with the leaflet package (v. 2.2.1) using GeoJSON data for state ZIP-code boundaries publicly available. B. Median geographic distances in Kilometers, highlighting variability in spatial distributions within and across communities.

**Figure S5:**
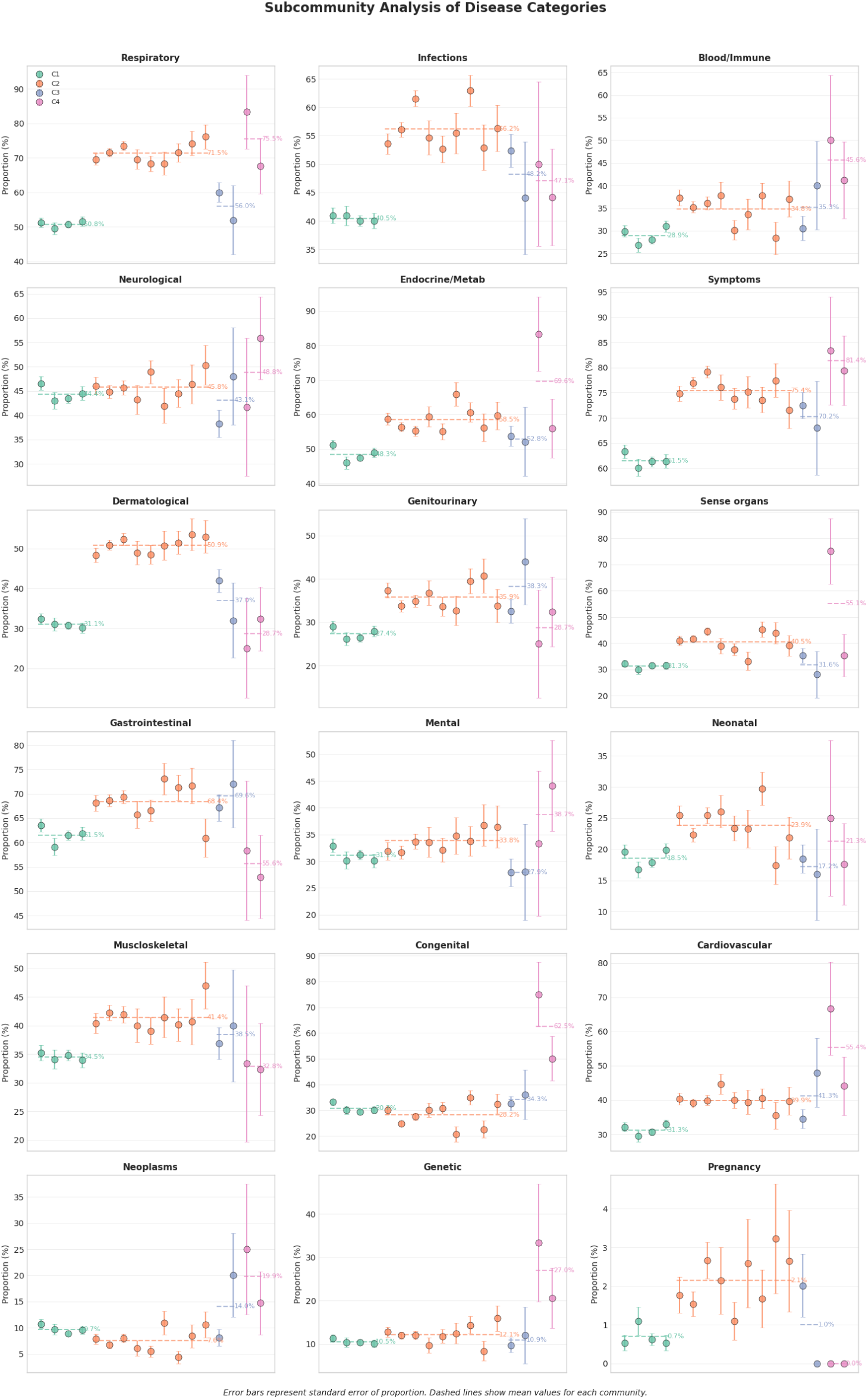
Proportion of phenotypes at the subcommunity level. CI95 for all proportions are shown and colored by community C1, C2, C3 and C4. The proportion is defined as the number of individuals in each community that reported, in at least one visit, a health condition in the corresponding category over the total. Error bars indicate 95% confidence intervals for proportions calculated using the Wilson score method.

**Figure S6:**
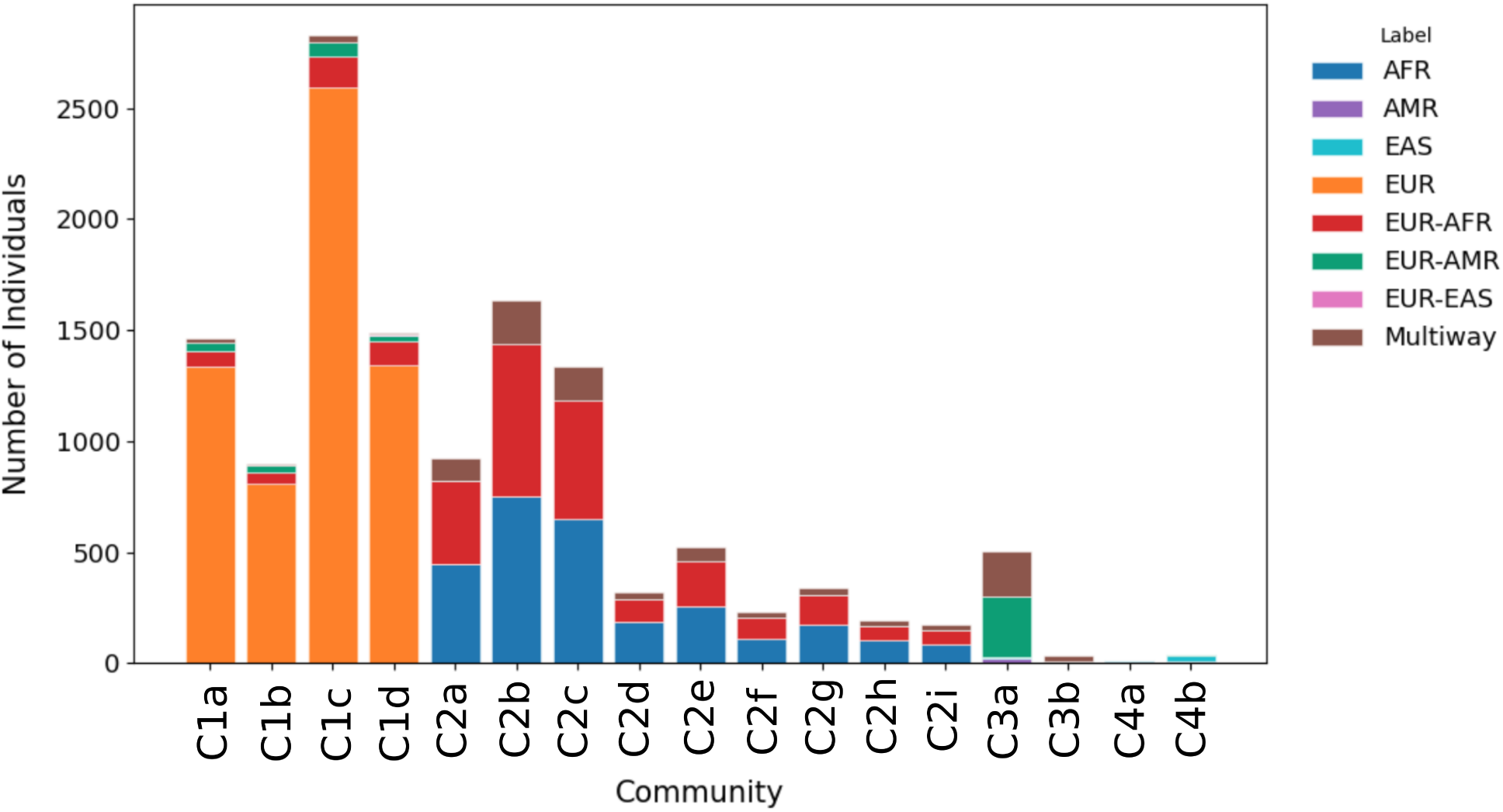
Number of BIG individuals in each subcommunity. We show the number of BIG individuals in each subcommunity , from the total of 13,143 incorporated. Inferred ancestries categories proportion by subcommunity are shown.

**Figure S7:**
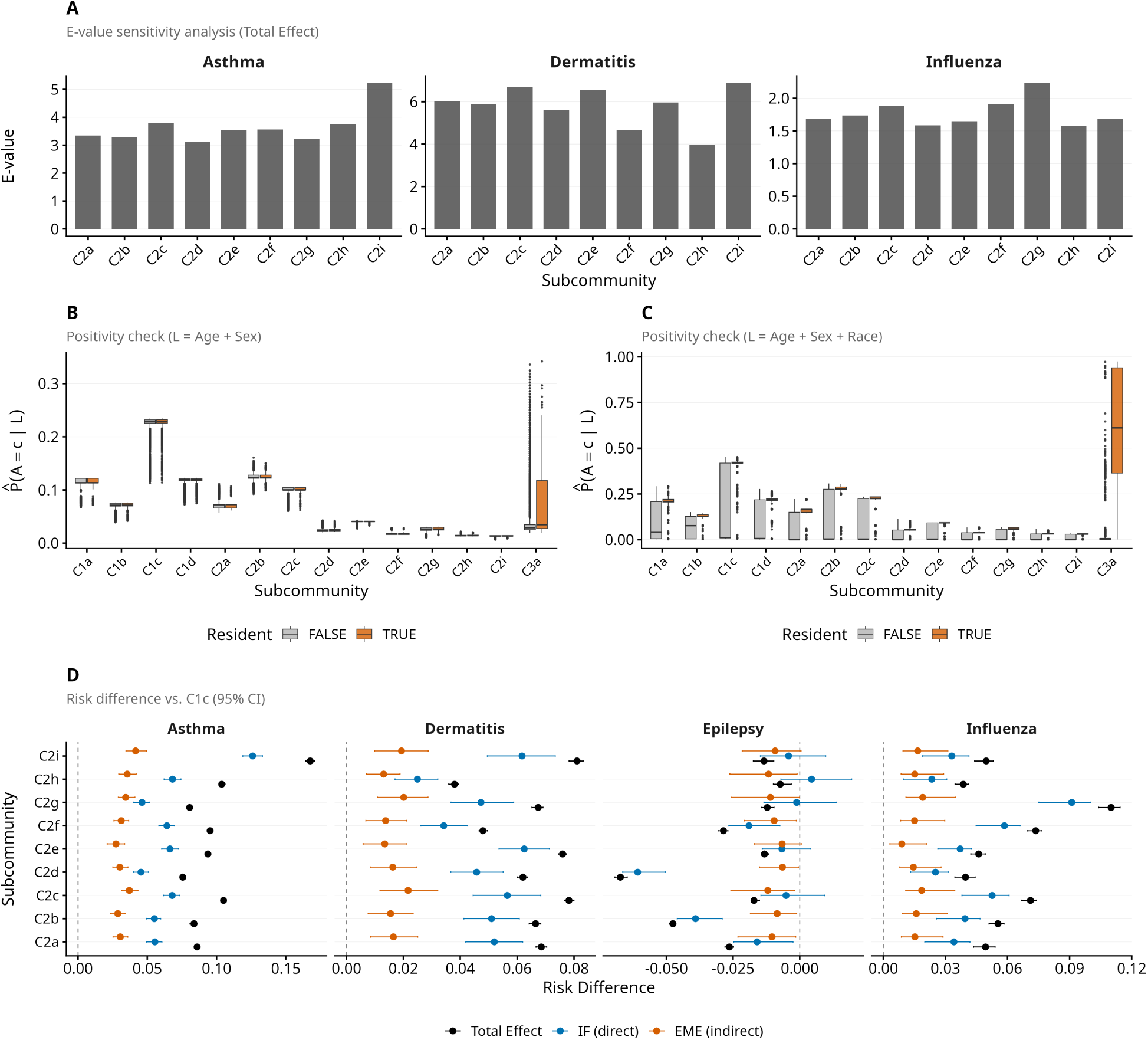
Sensitivity analyses and effect decomposition for the interventional mediation analysis. (**A**) E-value sensitivity analysis for total effects. E-values quantify the minimum confounder-exposure and confounder-outcome association strength needed to explain away observed effects. Asthma and dermatitis show E-values of 2–7, indicating substantial unmeasured confounding would be required to nullify these associations; influenza effects are more modest (E-values 1.5–2.2). (**B–C**) Positivity diagnostics showing propensity score distributions for each subcommunity. Orange: actual residents; grey: non-residents. Panel B conditions on age and sex; Panel C adds race. Adequate overlap indicates positivity is satisfied; overlap decreases with race adjustment, and C3 shows poor overlap. (**D**) Interventional mediation decomposition adding self-reported race as confounder. C2 subcommunities are shown relative to C1c: total effect (TE, black), independent effect (IF, blue), and environmental mediated effect (EME, orange). Error bars: 95% bootstrap CIs. EME captures effects mediated through PM_2.5_ and poverty; IF represents residual effects through other pathways.

**Figure S8:**
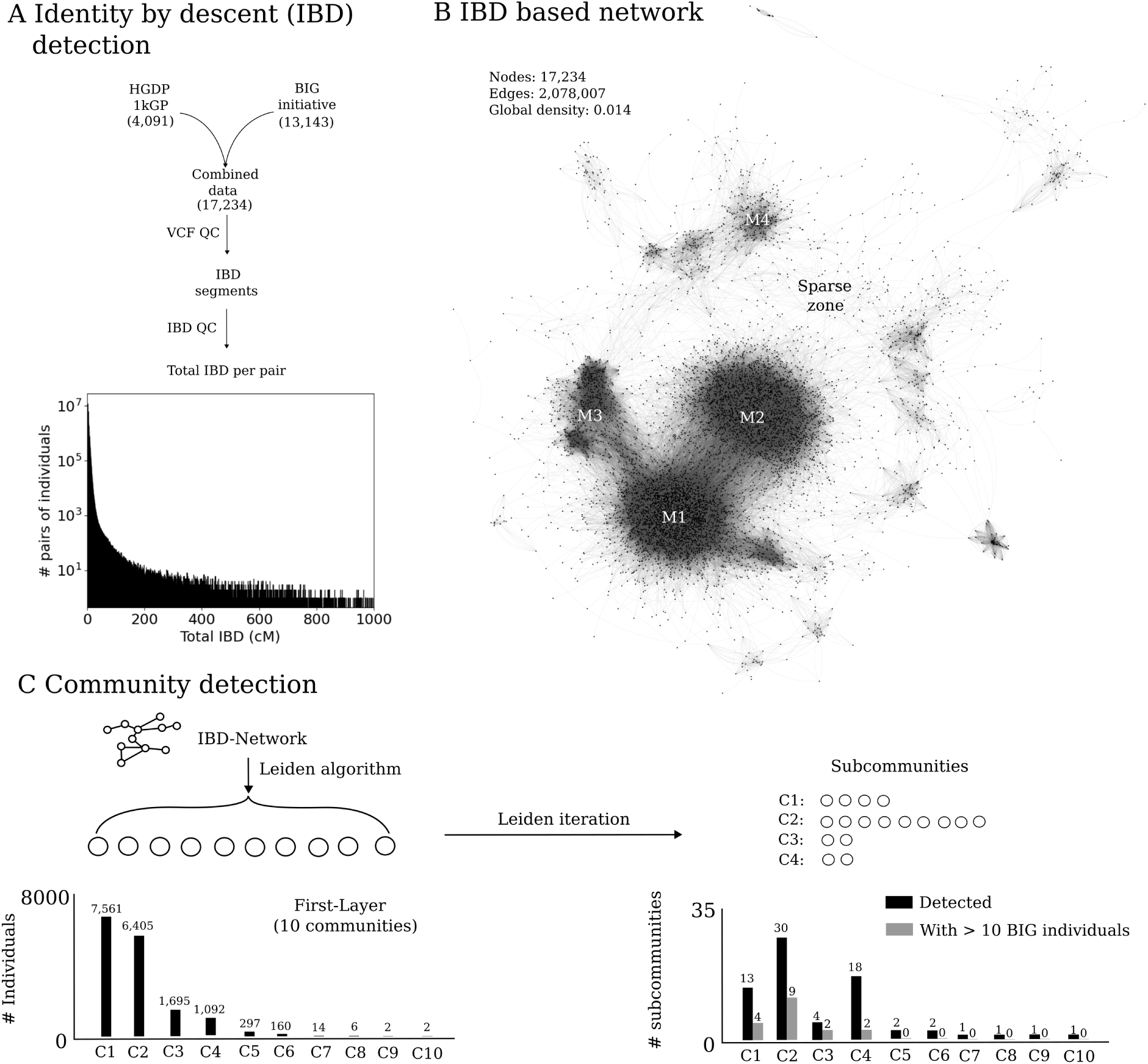
Distribution of identity-by-descent (IBD) values and corresponding network analysis. (A) Histogram showing the exponential decay of total IBD (cM). (B) IBD-based network with 17,234 nodes, revealing four dense communities (M1 to M4) and sparse zones. Fruchterman-Reingold layout was used to do the plot. (C) Bar plot of community sizes after the first Leiden iteration, with C1 to C4 containing over 1,000 individuals. (D) Number of subcommunities distribution, showing no direct correlation with parent community sizes.

**Table S1:** Standardization of self-reported race categories in electronic health records. We harmonized diverse race classifications to ensure analytical consistency and precision, following HL7 v2.5 standard terminologies while eliminating potentially problematic or outdated entries. This standardization process maintains demographic insights while supporting rigorous statistical analysis across our cohort (Popejoy, 2021). Legacy free-text entries such as “Caucasian” are preserved here as recorded in the EHR; their inclusion reflects historical clinical record-keeping and not endorsement of the term, and they are harmonized to the standardized HL7 v2.5 “White” category for analysis.

| Self-Reported Race | Number of individuals in self-reported race | Standardized HL7 v2.5 Race Category | Number of individuals in Standardized HL7 v2.5 Race Category |
| --- | --- | --- | --- |
| Black or African American | 4905 | Black or African American | 5279 |
| African American | 392 |  |  |
| White | 5672 | White | 6115 |
| White or Caucasian | 89 |  |  |
| Caucasian | 354 |  |  |
| Amer. Indian/Alaska Native | 8 | American Indian or Alaska Native | 35 |
| Native American | 27 |  |  |
| Asian | 68 | Asian | 69 |
| Asian or Pacific Islander | 1 |  |  |
| Native Hawaiian/Other Pacific Islander | 4 | Native Hawaiian or Other Pacific Islander | 6 |
| Pacific Islander | 2 |  |  |
| Other | 14 | Other race | 922 |
| Other/Unknown | 410 |  |  |
| Amharic (East African) | 1 |  |  |
| East Indian | 2 |  |  |
| Hispanic | 268 |  |  |
| Middle Eastern | 3 |  |  |
| Multiple | 224 | Unknown race | 45 |
| Unavailable | 37 |  |  |
| Declined | 4 |  |  |
| Patient Declined to Answer | 3 |  |  |

